# Transcriptional targets of senataxin and E2 promoter binding factors are associated with neuro-degenerative pathways during increased autophagic flux

**DOI:** 10.1101/2022.04.06.486307

**Authors:** Aaron E. Casey, Wenjun Liu, Leanne K. Hein, Timothy J. Sargeant, Stephen M. Pederson, Ville-Petteri Mäkinen

## Abstract

Autophagy is an intracellular recycling process that degrades harmful molecules, maintains optimal composition of cellular organelles and enables survival during starvation. Previous studies have demonstrated how transcription factors (TFs) can increase autophagy with therapeutic potential (impaired autophagy in the ageing brain, in particular, may be an important risk factor for dementia). To investigate the transcriptional regulation of autophagy from a systems perspective, we induced autophagy by amino acid starvation and mTOR inhibition in HeLa, HEK 293 and SH-SY5Y cells and used RNA-seq to measure gene expression at three time points. We observed 453 differentially expressed (DE) genes due to starvation and 284 genes due to mTOR inhibition (P_FDR_ < 0.05 in every cell line). Pathway analyses confirmed enrichment of genes implicated in Alzheimer’s (P_FDR_ < 0.001 in SH-SY5Y and HeLa) and Parkinson’s (P_FDR_ ≤ 0.024 in SH-SY5Y and HeLa) diseases and amyotrophic lateral sclerosis (ALS, P_FDR_ < 0.05 in 4 of 6 experiments). We then integrated Signaling Pathway Impact Analysis and TF target enrichment testing to predict which TF target genes were contributing to pathway perturbation. Differential expression of the Senataxin (SETX) target gene set was predicted to activate multiple neurodegenerative pathways (P_FDR_ ≤ 0.04). Notably, SETX is a causal gene for a rare form of ALS. In the SH-SY5Y cells of neuronal origin, the E2F transcription family was predicted to activate Alzheimer’s disease pathway (P_FDR_ ≤ 0.0065). SETX and E2F may be important mediators of transcriptional regulation of autophagy and may provide new therapeutic opportunities for neuro-degenerative conditions.

## Introduction

Maintaining energy homeostasis is essential for cells and biological organisms to survive and thrive. Throughout most of human history, perturbations to energy metabolism were due to starvation that stunted growth and development [1,2], while in modern populations metabolic health is challenged by sedentary life style, excess adiposity and ageing [3–5]. There is evidence that both energy extremes involve the same cellular processes that maintain energy homeostasis [6–8] and that these disruptions may be important drivers for common diseases such as diabetes and cancer [1,9–11]. Autophagy is one such process: it is responsible for recycling cellular materials into energy resources during periods of nutrient deprivation [12–14], but it also has an important role in maintaining the optimal composition of cellular organelles during periods of abundance [15]. Importantly, autophagy is affected by ageing [16] and impaired autophagy in the ageing brain, in particular, may be an important risk factor for Alzheimer’s and Parkinson’s diseases [17–20]. For these reasons, our long-term goal is to understand how autophagy and energy metabolism are regulated in human cells and to use this new fundamental knowledge towards new treatments for age-associated diseases.

Previous studies on autophagy regulation have revealed multiple pathways and genes [21,22], of which mammalian target for rapamycin (mTOR) and transcription factor EB (TFEB) are the best characterized [23,24]. A specific sequence, the coordinated lysosomal expression and regulation (CLEAR) motif seems to be the preferred DNA binding target for TFEB and its transcription factor family [25] and it may represent a key mechanism by which external conditions (e.g. starvation) exert a cascade of adaptation through mTOR, TFEB and the promoters of downstream autophagy genes [23]. As the name implies, the CLEAR motif is present in the promoters of lysosomal genes. This is important because the lysosome is the end-terminal of autophagic cascades [26] and it is responsible for the final degradation and recycling of materials including the two Alzheimer proteins, tau and amyloid-beta, that accumulate in the brains of affected individuals [20,27,28]. Lastly, we and others have identified genetic associations between autophagy and dementia [17,29,30]. These findings motivated us to explore the transcriptional responses associated with starvation-induced increase in autophagy and to investigate potential links between these responses and neuro-degenerative processes.

The aim of this study was to characterize how the transcriptome changes in response to starvation or mTOR inhibition in model systems where we also see responses in autophagy. We used genetically engineered human cells where we could confirm the changes in autophagic flux into the lysosome; this sets the experiments apart from previous work. Furthermore, we applied RNA sequencing at multiple time points and three cell lines to achieve robust systems-level understanding of which genes are reproducibly affected. The multi-faceted study design makes our study different from previous RNA-seq profiling experiments. Across the different cell line/treatment combinations, we report unexpected associations with differentially expressed genes and autophagic flux and characterize universal expression patterns and their predicted driver genes that overlap with neuro-degenerative disease processes.

## Results

### Overview of transcriptome responses

We collected RNA-seq data at baseline and after two interventions (mTOR inhibition or amino-acid starvation) in three monoclonal cell lines (HeLa, HEK 293 and SH-SY5Y). Initially, 12 differential expression analyses were conducted (2 time points × 2 treatments × 3 cell lines = 12). The two time points were then combined to focus only on the most robust and consistent signals (details in Methods). The resulting six lists of differentially expressed (DE) genes were used for further analyses and we refer to them as the six DE “experiments” throughout the text (2 treatments × 3 cell lines = 6, Supplementary Figure S1, Supplementary Tables S1-S6).

A total of 16,506 genes were detectable in at least one cell line and 11,202 (67.9%) were detectable in every cell line (Figure 1A). We observed 8,914 DE genes due to starvation in at least one cell line, of which 456 (5.1%) were classified as DE genes in every cell line (Figure 1B). We also observed 6,226 DE genes due to mTOR inhibition in at least one cell line; 285 (4.6%) of these were classified as DE genes in every cell line (Figure 1C). We identified 5,672 DE genes associated with starvation or mTOR inhibition that were up-regulated in at least one cell line (Figure 1D, inconsistent DE genes that were significantly up-regulated in one cell line but significantly down-regulated in another were excluded). Of these, 1,541 (27.2%) genes were shared by both treatments. Lastly, we identified 5,543 down-regulated genes of which 1,741 (31.4%) were shared between treatments (Figure 1E).

**Figure 1:**
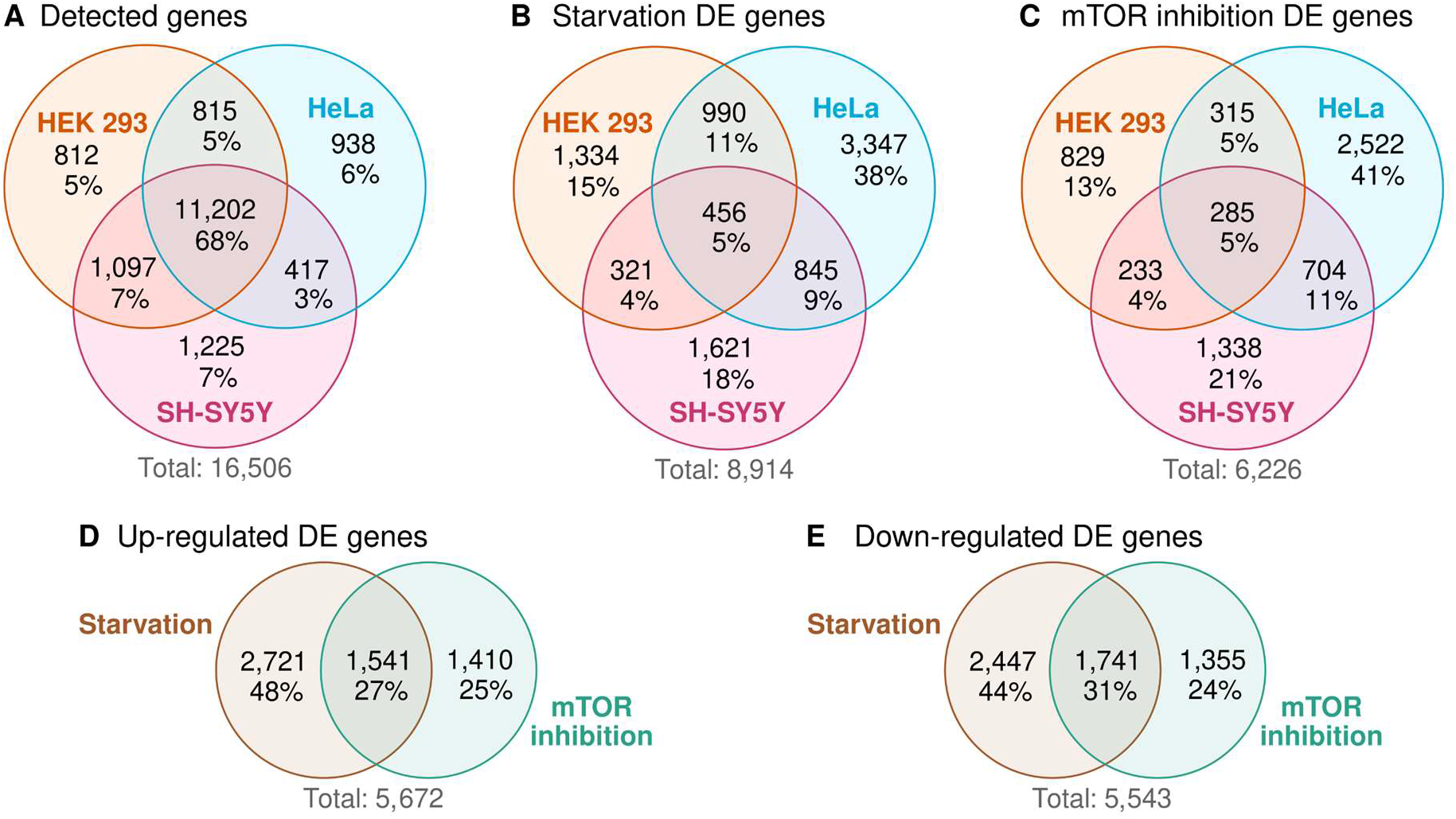
Overview of differentially expressed (DE) genes. **A**) Genes were considered detectable if there were >1.5 counts per million in >3 samples out of all samples from the same cell line. **B**) Genes that were DE between starved and control samples in at least one cell line. **C**) Genes that were DE between mTOR inhibited and control samples in at least one cell line. **D**) We collected DE genes associated with starvation or mTOR inhibition that were up-regulated in at least one cell line (inconsistent DE genes that were significantly up-regulated in one cell line but significantly down-regulated in another were excluded). **E**) Down-regulated DE genes associated with starvation or mTOR inhibition.

### Differentially expressed genes

The patterns of DE signals are summarized in Figure 2. P-values were adjusted for FDR as described in Methods (P_FDR_). For each plot, genes were ranked according to the maximum FDR rule: first, we determined the maximum P_FDR_ for each gene across the relevant experiments. Genes were then sorted according to maximum P_FDR_. Lastly, genes that were significantly up-regulated in one experiment, but down-regulated in another were excluded to maintain directional concordance.

**Figure 2:**
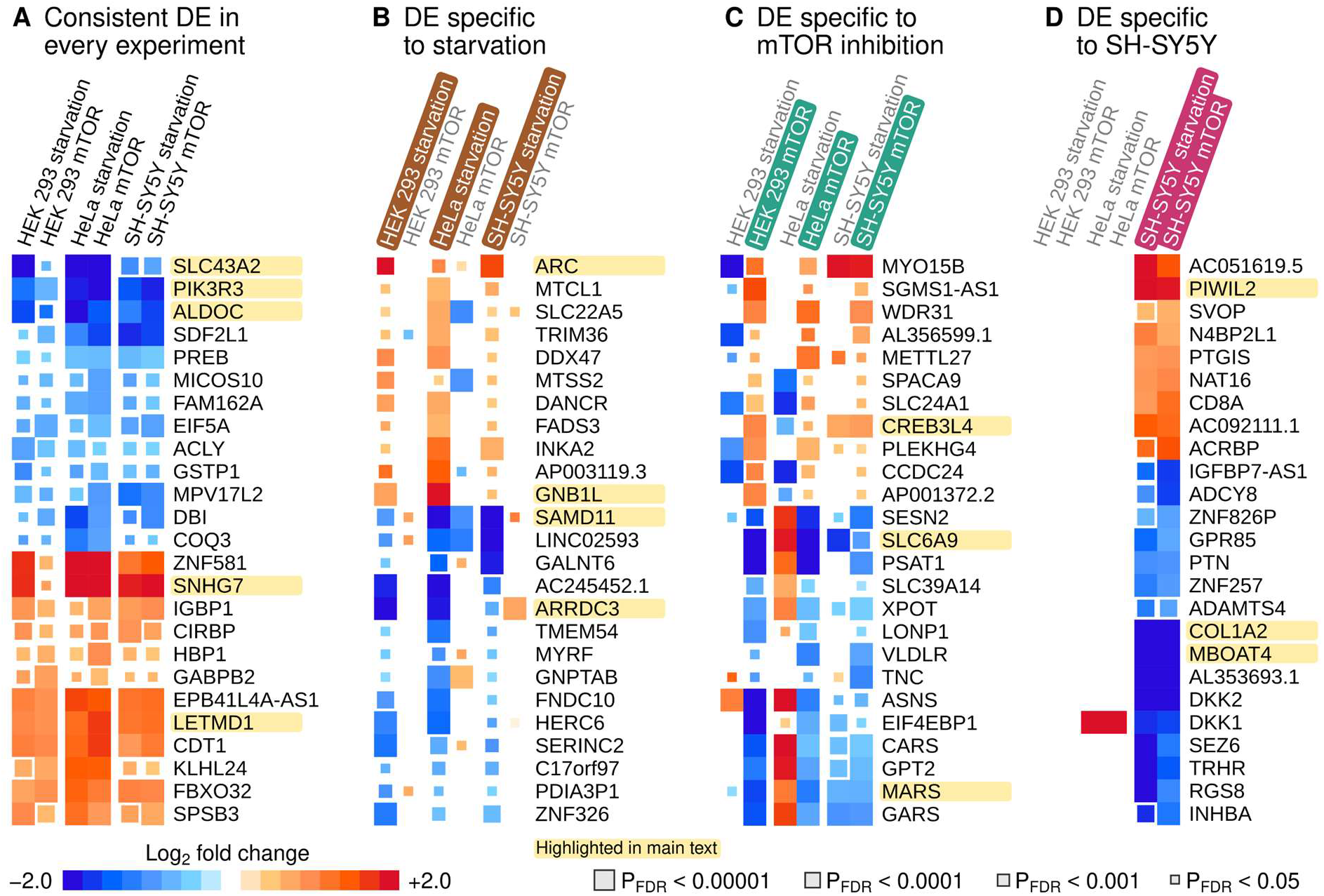
Top 25 differentially expressed (DE) genes based on the maximum FDR rule. Genes mentioned in the main text are highlighted for easier visual localization. **A**) Genes were sorted according to the maximum FDR-adjusted P-value across six experiments. Discordant genes that were significantly (P_FDR_ < 5%) up-regulated in one and down-regulated in another experiment were excluded. **B**) Genes were sorted according to the maximum P_FDR_ across all starvation experiments. We also required that all starvation responses were directionally concordant and that the mean log2 fold change across mTOR experiments was in the opposite direction. **C**) Genes were sorted according to the maximum P_FDR_ across all mTOR inhibition experiments. We required that all mTOR inhibition responses were directionally concordant and that the mean log2 fold change across starvation experiments was in the opposite direction. **D**) Genes were sorted according to the maximum P_FDR_ across responses in the SH-SY5Y cells. Missing signals were set to zero log2 fold change in other cell lines. We also required that the mean log2 fold changes in other cells were in the opposite direction to SH-SY5Y responses.

The gene with the lowest maximum P-value across all experiments (Figure 2A) was LETM1 Domain Containing 1 (LETMD1, P_FDR_ ≤ 5.0 × 10^-7^, mean logFC = 1.3, involved in phagocytosis). The greatest increase in relative expression was observed for a cancer-associated lncRNA that may inhibit autophagy (Small Nucleolar RNA Host Gene 7 or SNHG7, P_FDR_ ≤ 0.0041, mean logFC = 2.0). Genes that were down-regulated included a member of the PI3K family (Phosphoinositide-3-Kinase Regulatory Subunit 3, PIK3R3, P_FDR_ ≤ 5.6 × 10^-6^, mean logFC = −1.6), an L-amino acid transporter (Solute Carrier Family 43 Member 2, SLC43A2, P_FDR_ ≤ 0.0029, mean logFC = −1.7) and Aldolase Fructose-Bisphosphate C (ALDOC, P_FDR_ ≤ 0.00015, mean logFC = −1.6, a glycolysis gene associated with Alzheimer’s disease [31].

Top 25 genes ranked according to starvation response are shown in Figure 2B. The smallest maximum P-value across starvation experiments was observed for Activity Regulated Cytoskeleton Associated Protein (ARC, P_FDR_ ≤ 0.00074, mean logFC = 1.6, associated with memory and cognitive disorders [32]). The greatest increase in expression was observed for G Protein Subunit Beta 1 Like (GNB1L, P_FDR_ ≤ 0.0074, mean logFC = 1.1, associated with neurological disorders [33]). The top down-regulated gene was a transcriptional co-repressor involved in photoreceptor degradation and possibly autism (Sterile Alpha Motif Domain Containing 11, SAMD11, P_FDR_ ≤ 8.3 × 10^-5^, mean logFC = −1.9 [34]). Arrestin Domain Containing 3 (ARRDC3, P_FDR_ ≤ 0.00097, mean logFC = −1.8) was also down-regulated and it is involved in endocytic recycling [35] and lysosomal degradation of receptors [36].

Genes that were specifically affected by mTOR inhibition are shown in Figure 2C. Upregulated genes included CAMP Responsive Element Binding Protein 3 Like 4 (CREB3L4, FDR ≤ 0.0025, mean logFC = 0.86, a transcription factor involved in glucose and lipid metabolism). Of the down-regulated genes, Methionyl-TRNA Synthetase 1 (MARS) had the smallest FDR (FDR ≤ 1.3 × 10^-8^, mean logFC = −1.2, involved in alveolar disease). The greatest decrease in expression was observed for Solute Carrier Family 6 Member 9 (SLC6A9, FDR ≤ 3.1 × 10^-5^, mean logFC = −2.0) which is a glycine transporter associated with Alzheimer’s disease [37]. Of note, we observed an mTOR-specific pattern among the top 25 in HEK 293 and HeLa cells, but not in SH-SY5Y.

We isolated responses specific to SH-SY5Y in Figure 2D by applying the maximum FDR rule to the two SH-SY5Y experiments only (see details in figure caption). Piwi Like RNA-Mediated Gene Silencing 2 (PIWIL2, a piRNA regulator of autophagy and apoptosis [38]) exhibited the smallest maximum P-value (P_FDR_ ≤ 4.4 × 10^-6^, mean logFC = 2.1). The top-ranked down-regulated gene was Collagen Type I Alpha 2 Chain (COL1A2, P_FDR_ ≤ 1.4 × 10^-10^, mean logFC = −2.9, a structural component of collagen). Membrane Bound O-Acyltransferase Domain Containing 4 (MBOAT4), which stimulates autophagy [39], was also among the most down-regulated genes (P_FDR_ ≤ 5.2 × 10^-10^, mean logFC = −2.8).

### Canonical pathways enriched for differentially expressed genes

In the previous section, we observed multiple genes associated with autophagy and neuro-degeneration, however, a single DE gene in isolation does not reveal the wider biological consequences. To gain more robust insight into the biological impact, we investigated i) if genes from a known biological pathway were over-represented among DE genes and ii) if DE genes were likely to perturb a known pathway when considering the gene-gene interactions within the pathway. Selected results are depicted in Figure 3 and full statistics are available in Supplementary Tables S7-S18.

**Figure 3:**
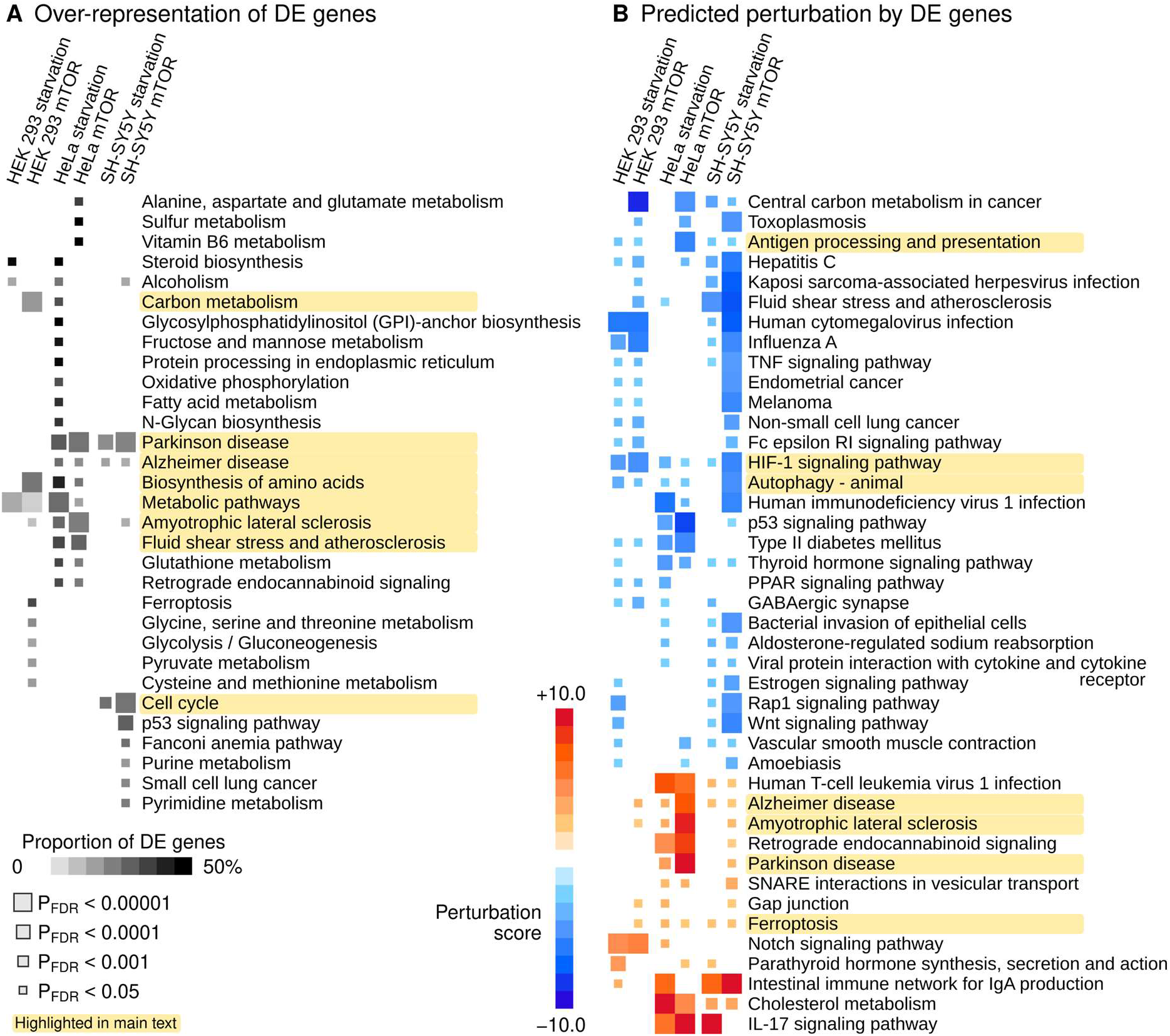
Enrichment of differentially expressed genes in the Kyoto Encyclopedia of Genes and Genomes pathway repository. **A**) Over-representation analysis of DE genes. Pathways that produced a significant signal (P_FDR_ < 0.05) in at least one experiment are shown. **B**) Normalized perturbation scores from Signaling Pathway Impact Analysis. A negative (positive) score implies that the aggregate impact of DE genes is likely to decrease (increase) the activity of a pathway. Pathways that were directionally concordant (all significant signals in the same direction) and that produced at least three significant signals (P_FDR_ < 0.05) are included.

We identified 31 KEGG pathways over-representing DE genes in at least one of the six experiments (Figure 3A). No pathway was significant in every experiment. The most consistent over-representation signals included Parkinson’s and Alzheimer’s diseases and Amyotrophic lateral sclerosis that were significant in four out of six experiments (highlighted in Figure 3A). Significant signals were also observed for Metabolic pathways, Cell cycle, Biosynthesis of amino acids, Carbon metabolism and Fluid shear stress and atherosclerosis.

Perturbation tests revealed multiple pathways that were likely to be activated or inhibited by the changes in gene expression (Figure 3B). We observed a directionally consistent activation of the KEGG Alzheimer’s disease pathway (perturbation z-scores between +2.15 and +7.63) and Ferroptosis (z-scores between +1.81 and +3.27) across all six experiments (highlighted in Figure 3B). We also observed unexpected but consistent association for inhibition of the Autophagy pathway (z-scores between −5.3 and −1.58). HIF-1 signaling was predicted to be inhibited across all experiments. Multiple immune-system pathways such as Antigen processing and presentation were also predicted to be inhibited due to DE.

### Transcription factor target gene sets enriched for differential expression

Genome-wide regulatory processes are not yet fully understood and may be missed by existing pathway definitions. For this reason, we repeated the over-representation analysis from the previous section but replaced the KEGG pathways with transcription factor target (TFT) gene sets (defined in Methods).

We observed 30 TFT gene sets that over-represented DE genes in every experiment. Based on the literature, TFEB may be a key regulator of autophagy [23] and, as expected, the predicted TFEB targets were over-representing DE genes associated with autophagy also in our study (highlighted in Figure 4A). Noteworthy signals related to neurological health include the MORC Family CW-Type Zinc Finger 2 set (*MORC2* is associated with multiple neurological conditions), the Senataxin set (*SETX*, also known as Amyotrophic lateral sclerosis 4 protein), the THAP Domain Containing 1 set (*THAP1* is associated with the neurodevelopmental disease dystonia 6) and SPT16 Homolog set (*SUPT16H* is associated with neurodevelopmental problems).

**Figure 4:**
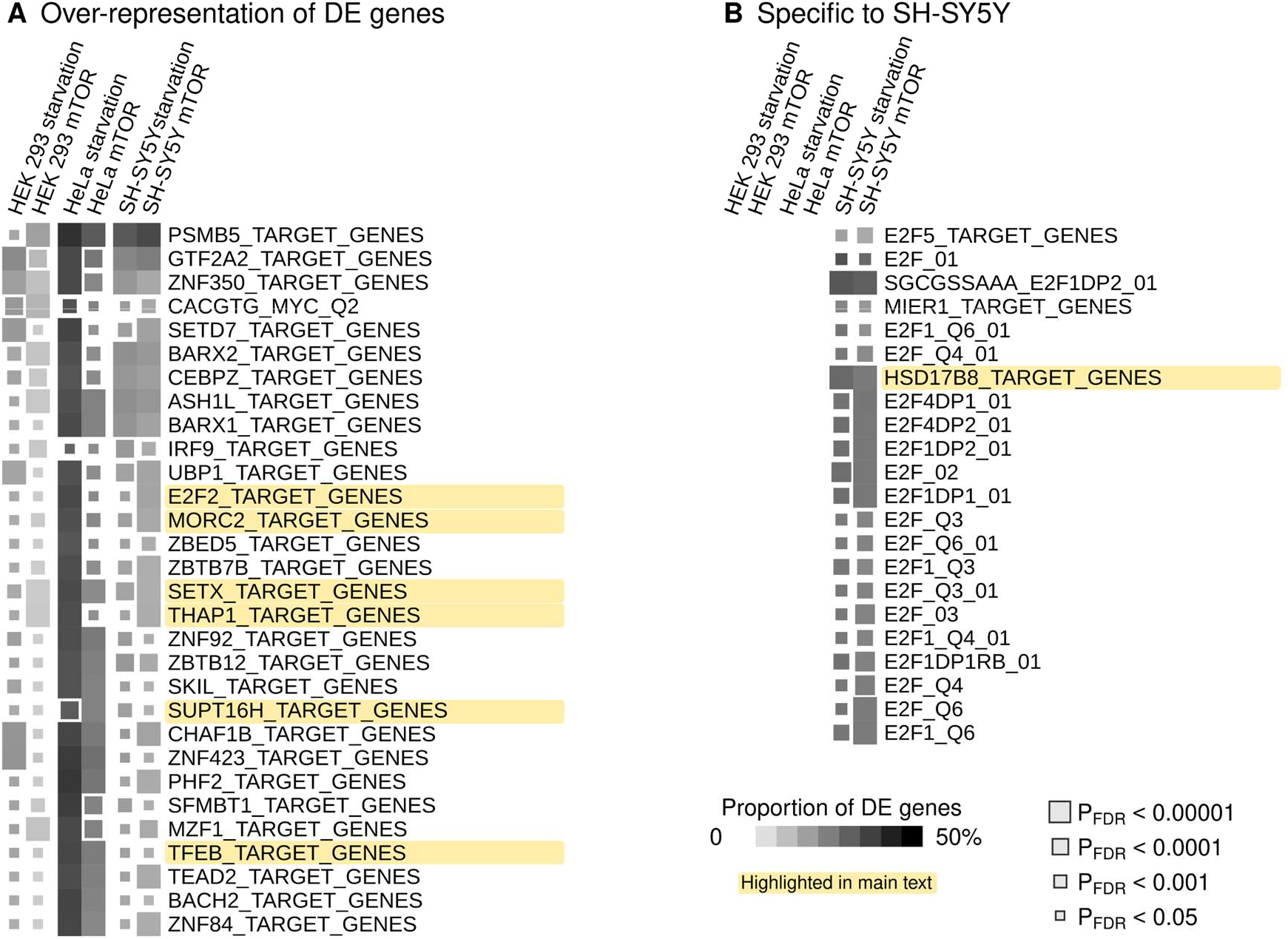
Enrichment of differentially expressed genes in transcription factor target (TFT) sets. **A**) Over-representation analysis of DE genes. Pathways that produced a significant signal (P_FDR_ < 0.05) in at least one experiment are shown. **B**) DE enrichment within TFT sets in SH-SY5Y cells but not in other cells.

We identified 22 gene sets that were enriched for DE genes in both SH-SY5Y experiments but not in other cells (Figure 4B); 20 belonged to the same E2F family that shared most of their target genes (note also *E2F2* in Figure 4A). The strongest signal was observed for the Hydroxysteroid 17-beta Dehydrogenase 8 (*HSD17B8*) gene set (P_FDR_ ≤ 7.9 × 10^-16^). Full results are available in Supplementary Tables S19-S24.

### Transcription factors as mediators between differential expression and canonical pathways

To compare the pathway and TFT responses, we first identified shared genes between a KEGG pathway and a TFT set, and then calculated the perturbations scores for this shared subset of genes. We observed ten pairs of transcription factors and KEGG pathways that satisfied P_FDR_ < 0.05 across every experiment – all of them were predicted to have increased activity due to the DE pattern and seven of them were pairings with KEGG neuro-degenerative diseases (Figure 5A, full results in Supplementary Tables S25-S30). Moreover, three signals involved the THAP1 gene set (paired with Alzheimer’s disease, ALS and viral infection) and four involved the SETX gene set (paired with Alzheimer’s, Parkinson’s, Huntington’s and ALS).

**Figure 5:**
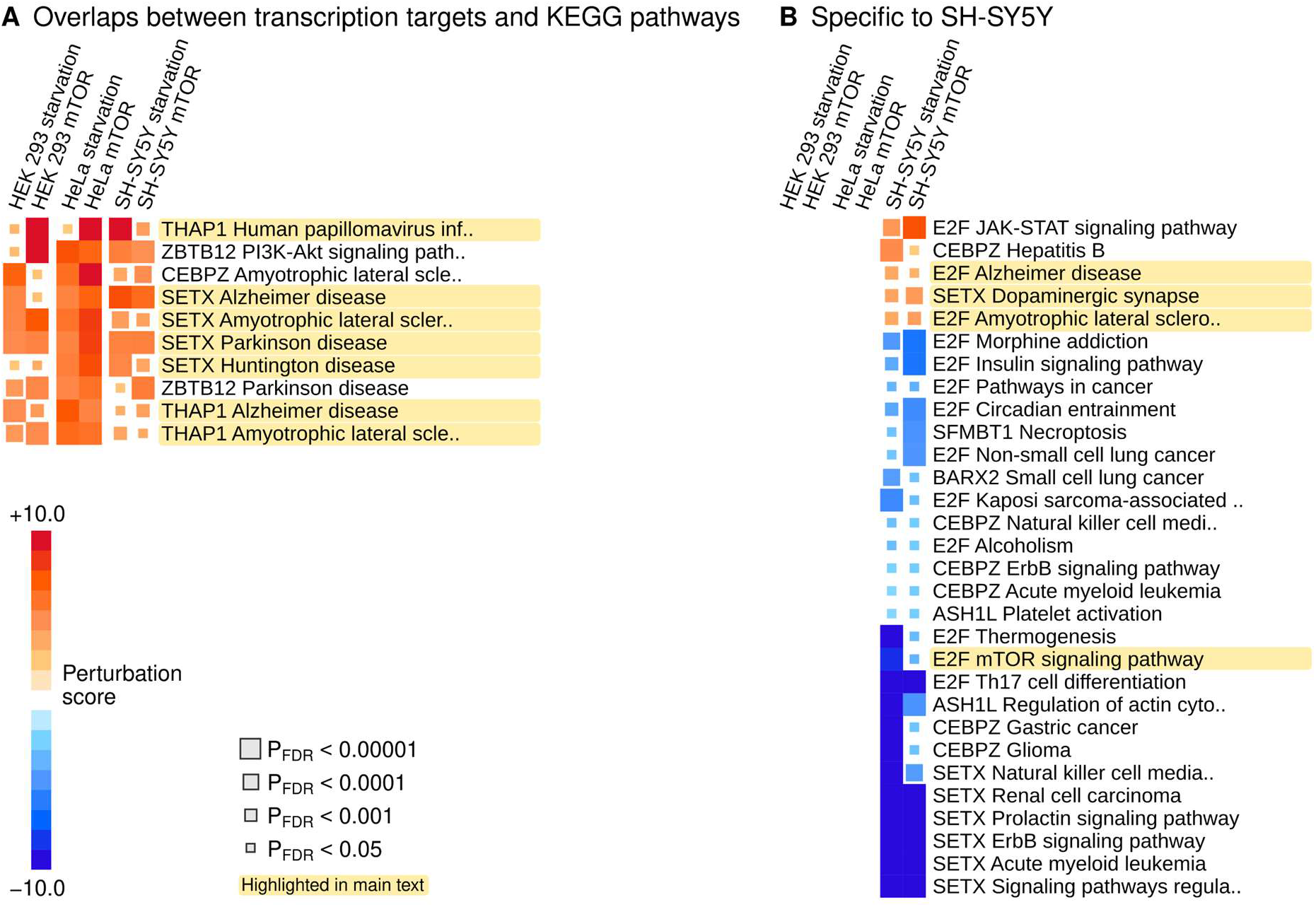
Combined perturbation analysis of canonical pathways and TFT sets. First, we identified DE genes that were shared between a KEGG pathway and TFT sets. Then, we used Signaling Pathway Impact analyses to test if the shared genes would impact the activity of the KEGG pathway. Therefore, the perturbation scores are predictions on the potential regulatory effects differentially expressed transcription factor target genes will have on canonical pathways. **A**) TFT-pathway pairs that showed directionally consistent and significant (P_FDR_ < 0.05) perturbation scores across every experiment. **B**) TFT-pathway pairs that showed directionally consistent and significant (P_FDR_ < 0.05) perturbation scores in the two experiments on SH-SY5Y cells but no significant signals in other cells.

We also found 32 pairs of TFT sets and pathways that were specific to SH-SY5Y (Figure 5B). Notably, the perturbation scores were mostly negative, which suggests that the DE of the TFT sets may result in the inhibition of these pathways. Exceptions included SETX and Dopaminergic synapse (P_FDR_ ≤ 0.00021), and multiple pathways perturbed by the E2F family of transcription factors, such as E2F and Alzheimer’s disease (P_FDR_ ≤ 0.0065).

## Discussion

To investigate the transcriptional regulation of autophagy in living human cells, we induced autophagic flux by amino acid starvation and mTOR inhibition and investigated the transcriptional responses in HeLa, HEK 293 and SH-SY5Y cell lines. Increase in autophagic flux was confirmed by a tandem-fluorescent LC3 assay (tf-LC3) and gene expression was quantified by RNA sequencing. We found that the KEGG autophagy pathway was inhibited at 15 h and 30 h after treatment, while pathways associated with neuro-generative diseases were activated. In particular, our results suggest that transcription target genes assigned to SETX and E2F may represent important regulatory mediators that connect energy metabolism, autophagy and cellular stress with Alzheimer’s and Parkinson’s diseases.

### Predicted inhibition of autophagy pathway

Previous literature and the tf-LC3 assays in this study show how autophagic flux increased in response to starvation or mTOR inhibition [14,24,40]. Against our expectations, we observed significant inhibition of the KEGG Autophagy pathway and possible autophagy regulators such as *ARRDC3* and *PIKR3* in the RNA-seq data. Moreover, putative autophagy inhibitors *LETMD1* and *SNHG7* were up-regulated. We also checked an earlier small pilot study to be sure that the treatment groups were not mislabeled (i.e. wrong sign of the statistical signal), but we observed matching direction of DE (data not shown).

We chose the time points of 15 h and 30 h based on time-series experiments to capture the inflection and saturation points of the autophagy response. A recent report on the dynamics of autophagy suggests that autophagy responses with respect to vesicular flux start within 10 min of treatment and saturate by 15 h [41]. In the first phase, mTOR Complex 1 inhibition by rapamycin induces an increase in autophagosomes which represents the initial packaging of molecular cargo into vesicles. Next, the autophagosomes fuse with lysosomes to form autolysosomes. Lastly, the autolysosomes degrade and the contents are recycled. The authors found that these three stages reached a steady state by 15 h where the numbers of autophagosomes and autolysosomes stabilize. Other studies have also demonstrated autophagic flux is still supported at late timepoints such as 8 h and 24 h in mouse embryonic fibroblasts [42], and 48 h in HeLa cells, as demonstrated by measurement of LC3-II with and without a lysosomal inhibitor drug [43]. Our results from the tf-LC3 assay, which tracks the proportion of LC3 within acidic autolysosomes, are compatible with these findings, although we observed stabilization at 30 h rather than 15 h in most cases.

Extrapolation of the dynamic autophagy process to transcriptional regulation may explain the negative DE we observed. We did not see substantial changes in gene expression at 1 h, which means that there was limited if any immediate transcriptional response associated with the initial increase in autophagosomes. On the other hand, by 15 h the transcriptome was responding to the nutrient deprivation, while the tf-LC3 assay was starting to level off. It is plausible that expressing autophagy genes at 15 h onward may become less of a priority for the cell and the relative expression of the pathway is subsequently decreased.

### Senataxin

SETX was first discovered via ataxia-associated mutations in a human homolog of the yeast gene Sen1 [44] and numerous additional mutations have since been reported that are associated with a rare type of ALS [45,46]. Initial studies showed that SETX helps to remove unintended DNA-RNA hybrid molecules (R-loops) that would otherwise promote genomic instability [47,48]. Interestingly, recent evidence indicates that SETX may be an important regulator of autophagy, especially with respect to the removal of stress granules that form when a cell is starved or under other types of environmental pressure [47]. For example, Richard et al. investigated a SETX knock-out [49] and reported that “SETX depletion inhibits the progression of autophagy, leading to an accumulation of ubiquitinated proteins, decreased ability to clear protein aggregates, as well as mitochondrial defects” which describes most neuro-degenerative diseases with features of proteinopathy. In another study, Bennet et al. induced SETX over-expression that disrupted the cell cycle of HEK 293 cells and they concluded that neurons due to their long RNA transcripts (i.e. propensity for R-loops) may be particularly vulnerable if SETX expression is outside the optimal range [50].

We observed up-regulation of genes that were implicated in Alzheimer’s and Parkinson’s disease, respectively, and predicted to be downstream targets of SETX (Figure 5A). On the other hand, SETX itself was not differentially expressed, which could be the result of tightly controlled expression range or transient expression patterns that are characteristic of transcription factors. Given the generic nature of our transcriptome findings, further studies of SETX may benefit from expanding the focus from ataxia and ALS to other types of neuro-degenerative diseases and focusing on energy restricted cellular milieu that may be characteristic to an ageing brain.

### E2 promoter binding factors

The E2F family of transcription factors is implicated in the regulation of energy metabolism, adipose tissue, obesity and growth in general [51–54] and our results from starved and mTOR-inhibited cells fit this picture well. The first family member, E2F1, is the most extensively studied. E2F1 binds with retinoblastoma protein to induce autophagy in cancer cells [54] and, inversely, E2F1 knockout inhibits autophagy to increase brown fat formation [52]. In Drosophila, E2F1 enables the regulation of TOR Complex 1 independent of insulin or amino acid pathways [53] and interacts with the cell cycle in a biphasic manner to promote organismal growth [51]. The E2F1 protein may be up-regulated in people with Down’s syndrome and amyloid-beta deposition [55].

The E2F signals we observed are most likely explained as universal consequences of energy restriction across cell types. In the neuroblastoma cell line, the E2F-targeted portions of cancer pathways, thermogenesis, mTOR signaling, insulin signaling, and circadian entrainment were all inhibited (Figure 5B), as one would expect based on the previous research on E2F1. On the other hand, ALS and Alzheimer’s disease pathways were predicted to be activated. Our data cannot reveal causal relationships, but we speculate that E2F transcription factors respond to age-associated metabolic dysfunction and may subsequently trigger neuronal apoptosis [56,57]. There is evidence that inducing E2F1 and E2F2 may help maintain genomic stability in neurons under toxic conditions [58] while other experiments showed that reducing E2F1 in mice improved the survival of dopaminergic neurons [57]. Given these complex and contradictory findings, additional research into the exact roles of each E2F family member in relation to human tissues is warranted.

### Strengths and weaknesses

The inclusion of three cell lines, three time points and two conditions provide statistical and biological robustness to our findings. HEK 293, HeLa and SH-SY5Y cells are established platforms for experimental studies and grow predictably in standard conditions, which helped us to maintain high consistency between cultures. On the other hand, these immortalized cells may differ substantially from human cells *in situ* and the interventions we chose are beyond the typical physiological stresses most cells would encounter. Hence, we caution against over-reaching conclusions about possible therapeutic targets among the top DE genes. Instead, these data should be interpreted as further evidence on the associations between energy metabolism, autophagy machinery and neuro-degenerative diseases, whereas the exact causal mechanisms may be highly dependent on the cell type or on an individual’s genetic profile.

The use of immortal cell lines allowed us to optimize monoclonal cultures that expressed the tf-LC3 construct. This was important to achieve a high signal-to-noise ratio for the fluorescence assay for autophagic flux. Furthermore, the technical quality and depth of the RNA-seq data were high and we used additional permutation tests to verify signals beyond the original pathway tools. For these reasons, we are confident that the analytical quality of the study is high.

### Conclusions

We conducted an experimental study to characterize transcriptomic changes associated with autophagy in three human cell lines. Our setup was not optimized for neuro-degenerative diseases beyond the neuronal SH-SY5Y cells, yet to our surprise we identified an enrichment of differentially expressed genes in Alzheimer’s and Parkinson’s disease pathways that emerged from the RNA-seq data. This reenforces the idea that autophagy and energy metabolism are intrinsically involved in these major human diseases. Furthermore, we identified senataxin and the E2F transcription factor family as potential mediators between transcriptional regulation of autophagy and neuro-degenerative conditions.

## Materials and methods

### Study design

The experimental part of the project comprised four subcomponents: i) an assay for autophagic flux, ii) the selection and culture of cell lines, iii) time series design for mTOR inhibition and starvation, and iv) final experiments for transcriptomic analysis. Firstly, we used a tandem fluorescent LC3 (tf-LC3) assay to measure autophagic flux. LC3 is a core protein component of the autophagosome membrane that is eventually incorporated into the lysosome at the end of the vesicular autophagy pipeline [40]. The tf-LC3, which comprises the LC3 protein fused to a red fluorescence protein and a pH-sensitive green fluorescence protein, is incorporated into the autophagosome in the same manner as the native LC3. Once the tf-LC3 proteins reach the acidic interior of the lysosome the pH-sensitive green fluorophore is quenched while the red fluorophore is unaffected. Therefore, the ratio between red and green fluorescence indicates the proportion of tf-LC3 in lysosomes versus total cellular tf-LC3, which we use as a proxy for autophagic flux.

Secondly, we selected three different cell lines to identify consistent and universal RNA expression changes associated with changes in autophagy flux. We chose Hela and HEK 293 cells due to their robust growth in cultures and the SH-SY5Y due to their brain-tissue origin. Each cell line of interest was transfected with lentiviral particles that contained the sequence for the tf-LC3 construct under the cytomegalovirus promoter [59]. Multiple monoclonal lines were cultured for each cell line, each of which had total red and green fluorescence quantified by flow cytometry, thus allowing for selection of clones most appropriate for quantifying autophagic flux in the proposed experiments.

Thirdly, we subjected the clones to mTOR inhibition using 1 μM of AZD8055 (Selleck Chemicals LLC, Houston TX, USA) and to amino-acid starvation using Earl’s balanced salt solution (EBSS; MSD, Kenilworth NJ, USA) to induce autophagy. Temporal curves of autophagic flux were determined by measuring the tf-LC3 red/green ratio at 1 h intervals (Supplementary Figure S2). Based on the curves, we chose 1 h, 15 h and 30 h time points as the initial response, inflection point and saturation point of autophagic flux respectively. Three technical replicates were collected from every experimental arm. We observed no difference between the baseline and 1 h RNA profiles, thus only 15 h and 30 h time points were used for statistical analyses.

### Cell culture and materials

HeLa and HEK 293 cell lines were cultured in Dulbecco’s Modified Eagle Medium (DMEM; Life Technologies, Thermo Fisher Scientific, Waltham MA, USA), while SH-SY5Y cells were cultured in 1:1 DMEM:Ham’s F12 (MSD, Kenilworth NJ, USA). All three cell lines were maintained with 10% (v/v) foetal bovine serum (Life Technologies), and 5 mg/mL penicillin and streptomycin (MSD, Kenilworth NJ, USA) in a humidified atmosphere of 5% CO_2_ at 37°C. For RNA profiling, T_25_ flasks were seeded with 1.24 × 10^6^ cells from 80% confluent T_75_ flasks 24 h prior to the start of experiments.

Both the treated and non-treated samples were seeded from the same flask. Parental clones without the tf-LC3 proteins were used to calibrate the flow cytometer before measuring autophagic flux.

### RNA sequencing

Total RNA was extracted using the RNeasy Plus Mini Kit (QIAGEN, Hilden North Rhine-Westphalia, Germany) as per the manufacturer’s instructions (sample RNA concentration ≥15 ng/μL, ≤2809 ng/μL, median 361.5 ng/μL). The RNA library was prepared with indices and was sequenced on an Illumina NovaSeq 6000 S4 at 2 × 150 bp at the David R Gunn Genomics Suite in the South Australian Health and Medical Research Institute.

### RNA data processing

The scripts that were used for the analyses are available at https://github.com/Wenjun-Liu/Induced_autophagy. Default parameter settings were used at each step unless otherwise indicated. Each sample was sequenced to a median of 132 million paired reads per sample. We applied a three-step protocol to process raw reads into gene-level expression estimates. Firstly, we used cutadapt version 1.14 [60] to trim away low quality bases, adapters and other non-useful sequences. Secondly, trimmed reads were aligned to the human genome assembly GRCh38.p13 from Ensembl Release 98 [61] using STAR v2.7 [62]. Thirdly, total read counts for each gene (i.e. gene-level expression estimates) were quantified using featureCounts from the Subread package version 1.5.2 [63], with the setting 1 for fracOverlap and 10 for Q. Gene annotations were obtained from Ensembl Release 98 [61].

For each of the three steps, quality checks were performed using FastQC v0.11.7 (URL: https://www.bioinformatics.babraham.ac.uk/projects/fastqc/) and ngsReports [64]. We observed no issues related to low sequencing quality, variable GC content or high adapter content across the set of libraries. We considered a gene detectable for a cell line if we observed >1.5 counts per million in >3 samples out of 15, representing all samples from a complete treatment arm. A total of 16,506 (24.3%) out of 67,946 annotated genes were detectable in at least one cell line and 11,202 (16.5%) genes were detectable in every cell line. Lastly, we applied the conditional quantile normalization method to mitigate remaining artefacts from GC content and gene length in preparation for the statistical analysis [65].

### Differential expression analysis

We identified differentially expressed (DE) genes between the treated and untreated cell lines by quasilikelihood negative binomial generalised log-linear regression as implemented in edgeR [66,67]. We defined DE that exceeded the range of ±20% fold change as biologically meaningful [68]. We then used the quasi-likelihood F-test to calculate P-values. P-values were further adjusted by the Benjamini-Hochberg method of false discovery rates (P_FDR_) to account for multiple testing [11].

We conducted 12 initial DE analyses where we compared the expression levels at 15 h and 30 h against the baseline at 0 h (3 cell lines × 2 treatments × 2 time points, Supplementary Figure S1). Statistically significant genes (P_FDR_ < 0.05) were then selected for further investigation from each DE analysis. Given the overlap between significant genes at 15 h and 30 h time-points (mean 53.4% across cell lines and treatments), we included only those that showed significant DE in the same direction at both time points, as a strategy to focus on the most consistently changed genes. For a single estimate of foldchange, we used the mean log2 fold change across both time-points. Hence the final set of results comprised six separate DE listings (3 cell lines × 2 treatments × 1 combined time point, Supplementary Figure S1).

### Pathway enrichment analysis

We investigated i) if genes in pre-defined biological pathways were over-represented among DE genes and ii) to what extent DE genes were likely to perturb a given pathway when considering the known functional relationships between the pathway members. Firstly, over-representation of DE genes was tested with using goseq [12]. We included an offset term to account for bias due to gene size that can confound other over-representation approaches.

Secondly, we applied the Signaling Pathway Impact Analysis (SPIA) method to identify potentially perturbed pathways [69]. The SPIA adds to the results from goseq since it provides deeper functional insight into the consequences from altered gene expression. In SPIA, pathways are represented as networks of genes based on pathway topology and activating/inhibitory roles of individual genes. Perturbation is defined as the propagating effect from altering the expression of one or more genes within the network. Crucially, the SPIA algorithm predicts the accumulated perturbation effect from multiple DE genes and summarizes the total effect as a single numerical score. This perturbation score is directional: a negative score indicates down-regulation of a pathway, whereas a positive score indicates up-regulation. In this study, we used a novel permutation procedure to calculate the statistical significance of the perturbation score (Supplementary Figure S3).

Canonical pathway definitions were obtained from the Kyoto Encyclopedia of Genes and Genomes (KEGG) database [70]. We retrieved 312 KEGG pathways and converted them into an SPIA-compatible network format using the tool graphite [71]. Over-representation and perturbation tests were applied to each of the six DE listings, respectively. The threshold for significant over-representation was set at 5% FDR. The same threshold was also applied to define significant perturbation.

### Transcription factor target genes

We retrieved 957 transcription factor (TFT) gene sets from the Molecular Signatures Database version 7.2 [72,73] where, for a specific transcription factor, the TFT gene set was determined according to the binding sites or promoter binding motifs in the target genes. TFT gene sets were analysed for overrepresentation of DE genes the same way as the KEGG pathways, however, as the TFT definitions do not include interaction information between the target genes, we developed a strategy to combine the TFT information with KEGG pathway topologies. First, we determined subsets of genes that were shared between a KEGG pathway and a TFT gene-set; these subsets represent potential mechanisms by which a transcription factor may regulate a KEGG pathway. To test the regulatory potential further, we applied SPIA the same way as before, but using the subset of genes within the KEGG pathway (that were also TFT genes). KEGG pathways with P_FDR_ < 0.05 were considered to be significantly perturbed due to the given transcription factor.

## Author contributions

VPM and AC conceived and designed the study. AC, LH and TS managed laboratory resources and conducted cell experiments. WL and SP designed statistical experiments and methods, and analysed the data. AC, WL and VPM wrote the manuscript draft. All authors reviewed and edited the text.

## Data availability

RNA sequencing data with autophagic flux measurements will be available in the EBI ArrayExpress repository when the text is accepted for publication in a peer-reviewed journal. Analysis scripts are available in GitHub (URL: https://github.com/Wenjun-Liu/Induced_autophagy).

## References

[1] R. Du, R. Zheng, Y. Xu, Y. Zhu, X. Yu, M. Li, X. Tang, R. Hu, Q. Su, T. Wang, Z. Zhao, M. Xu, Y. Chen, L. Shi, Q. Wan, G. Chen, M. Dai, D. Zhang, Z. Gao, G. Wang, F. Shen, Z. Luo, Y. Qin, L. Chen, Y. Huo, Q. Li, Z. Ye, Y. Zhang, C. Liu, Y. Wang, S. Wu, T. Yang, H. Deng, L. Chen, J. Zhao, Y. Mu, D. Li, G. Qin, W. Wang, G. Ning, L. Yan, Y. Bi, J. Lu, Z.T. Bloomgarden, Early-Life Famine Exposure and Risk of Cardiovascular Diseases in Later Life: Findings From the REACTION Study, J. Am. Heart Assoc. 9 (2020). https://doi.org/10.1161/JAHA.119.014175.

[2] K. Hidayat, X. Du, B. Shi, L. Qin, Foetal and childhood exposure to famine and the risks of cardiometabolic conditions in adulthood: A systematic review and meta-analysis of observational studies, Obes. Rev. 21 (2020). https://doi.org/10.1111/obr.12981.

[3] B.T. Tam, J.A. Morais, S. Santosa, Obesity and ageing: Two sides of the same coin, Obes. Rev. 21 (2020). https://doi.org/10.1111/obr.12991.

[4] U.M. Kujala, V.-P. Mäkinen, I. Heinonen, P. Soininen, A.J. Kangas, T.H. Leskinen, P. Rahkila, P. Würtz, V. Kovanen, S. Cheng, S. Sipilä, M. Hirvensalo, R. Telama, T. Tammelin, M.J. Savolainen, A. Pouta, P.F. O’Reilly, P. Mäntyselkä, J. Viikari, M. Kähönen, T. Lehtimäki, P. Elliott, M.J. Vanhala, O.T. Raitakari, M.-R. Järvelin, J. Kaprio, H. Kainulainen, M. Ala-Korpela, Long-term leisure-time physical activity and serum metabolome, Circulation. 127 (2013) 340–348. https://doi.org/10.1161/CIRCULATIONAHA.112.105551.

[5] M. Juonala, C.G. Magnussen, G.S. Berenson, A. Venn, T.L. Burns, M.A. Sabin, S.R. Srinivasan, S.R. Daniels, P.H. Davis, W. Chen, C. Sun, M. Cheung, J.S.A. Viikari, T. Dwyer, O.T. Raitakari, Childhood Adiposity, Adult Adiposity, and Cardiovascular Risk Factors, N. Engl. J. Med. 365 (2011) 1876–1885. https://doi.org/10.1056/NEJMoa1010112.

[6] M.L. Steinhauser, B.A. Olenchock, J. O’Keefe, M. Lun, K.A. Pierce, H. Lee, L. Pantano, A. Klibanski, G.I. Shulman, C.B. Clish, P.K. Fazeli, The circulating metabolome of human starvation, JCI Insight. 3 (2018) 121434. https://doi.org/10.1172/jci.insight.121434.

[7] J.H. Lee, A. Park, K.-J. Oh, S.C. Lee, W.K. Kim, K.-H. Bae, The Role of Adipose Tissue Mitochondria: Regulation of Mitochondrial Function for the Treatment of Metabolic Diseases, Int. J. Mol. Sci. 20 (2019) E4924. https://doi.org/10.3390/ijms20194924.

[8] B.M. Spiegelman, J.S. Flier, Obesity and the Regulation of Energy Balance, Cell. 104 (2001) 531–543. https://doi.org/10.1016/S0092-8674(01)00240-9.

[9] C. García-Jiménez, M. Gutiérrez-Salmerón, A. Chocarro-Calvo, J.M. García-Martinez, A. Castaño, A. De la Vieja, From obesity to diabetes and cancer: epidemiological links and role of therapies, Br. J. Cancer. 114 (2016) 716–722. https://doi.org/10.1038/bjc.2016.37.

[10] R.K. Singh, P. Kumar, K. Mahalingam, Molecular genetics of human obesity: A comprehensive review, C. R. Biol. 340 (2017) 87–108. https://doi.org/10.1016/j.crvi.2016.11.007.

[11] P. Würtz, Q. Wang, A.J. Kangas, R.C. Richmond, J. Skarp, M. Tiainen, T. Tynkkynen, P. Soininen, A.S. Havulinna, M. Kaakinen, J.S. Viikari, M.J. Savolainen, M. Kähönen, T. Lehtimäki, S. Männistö, S. Blankenberg, T. Zeller, J. Laitinen, A. Pouta, P. Mäntyselkä, M. Vanhala, P. Elliott, K. H. Pietiläinen, S. Ripatti, V. Salomaa, O.T. Raitakari, M.-R. Järvelin, G.D. Smith, M. Ala-Korpela, Metabolic signatures of adiposity in young adults: Mendelian randomization analysis and effects of weight change, PLoS Med. 11 (2014) e1001765. https://doi.org/10.1371/journal.pmed.1001765.

[12] F. Pietrocola, Y. Demont, F. Castoldi, D. Enot, S. Durand, M. Semeraro, E.E. Baracco, J. Pol, J.M. Bravo-San Pedro, C. Bordenave, S. Levesque, J. Humeau, A. Chery, D. Métivier, F. Madeo, M.C. Maiuri, G. Kroemer, Metabolic effects of fasting on human and mouse blood in vivo, Autophagy. 13 (2017) 567–578. https://doi.org/10.1080/15548627.2016.1271513.

[13] R.C. Scott, O. Schuldiner, T.P. Neufeld, Role and Regulation of Starvation-Induced Autophagy in the Drosophila Fat Body, Dev. Cell. 7 (2004) 167–178. https://doi.org/10.1016/j.devcel.2004.07.009.

[14] S. Erbil-Bilir, D. Gozuacik, O. Kutlu, Autophagy as a Physiological Response of the Body to Starvation, in: V. Preedy, V.B. Patel (Eds.), Handb. Famine Starvation Nutr. Deprivation, Springer International Publishing, Cham, 2017: pp. 1–15. https://doi.org/10.1007/978-3-319-40007-5_69-1.

[15] A.L. Anding, E.H. Baehrecke, Cleaning House: Selective Autophagy of Organelles, Dev. Cell. 41 (2017) 10–22. https://doi.org/10.1016/j.devcel.2017.02.016.

[16] M.C. Barbosa, R.A. Grosso, C.M. Fader, Hallmarks of Aging: An Autophagic Perspective, Front. Endocrinol. 9 (2018) 790. https://doi.org/10.3389/fendo.2018.00790.

[17] S. Gao, A.E. Casey, T.J. Sargeant, V.-P. Mäkinen, Genetic variation within endolysosomal system is associated with late-onset Alzheimer’s disease, Brain J. Neurol. 141 (2018) 2711–2720. https://doi.org/10.1093/brain/awy197.

[18] D.A. Loeffler, Influence of Normal Aging on Brain Autophagy: A Complex Scenario, Front. Aging Neurosci. 11 (2019) 49. https://doi.org/10.3389/fnagi.2019.00049.

[19] A. Metaxakis, C. Ploumi, N. Tavernarakis, Autophagy in Age-Associated Neurodegeneration, Cells. 7 (2018) 37. https://doi.org/10.3390/cells7050037.

[20] L.S. Whyte, A.A. Lau, K.M. Hemsley, J.J. Hopwood, T.J. Sargeant, Endo-lysosomal and autophagic dysfunction: a driving factor in Alzheimer’s disease?, J. Neurochem. 140 (2017) 703–717. https://doi.org/10.1111/jnc.13935.

[21] I. Dikic, Z. Elazar, Mechanism and medical implications of mammalian autophagy, Nat. Rev. Mol. Cell Biol. 19 (2018) 349–364. https://doi.org/10.1038/s41580-018-0003-4.

[22] C. Di Malta, L. Cinque, C. Settembre, Transcriptional Regulation of Autophagy: Mechanisms and Diseases, Front. Cell Dev. Biol. 7 (2019) 114. https://doi.org/10.3389/fcell.2019.00114.

[23] C.J. Cortes, A.R. La Spada, TFEB dysregulation as a driver of autophagy dysfunction in neurodegenerative disease: Molecular mechanisms, cellular processes, and emerging therapeutic opportunities, Neurobiol. Dis. 122 (2019) 83–93. https://doi.org/10.1016/j.nbd.2018.05.012.

[24] Y. Wang, H. Zhang, Regulation of Autophagy by mTOR Signaling Pathway, in: Z.-H. Qin (Ed.), Autophagy Biol. Dis., Springer Singapore, Singapore, 2019: pp. 67–83. https://doi.org/10.1007/978-981-15-0602-4_3.

[25] M. Palmieri, S. Impey, H. Kang, A. di Ronza, C. Pelz, M. Sardiello, A. Ballabio, Characterization of the CLEAR network reveals an integrated control of cellular clearance pathways, Hum. Mol. Genet. 20 (2011) 3852–3866. https://doi.org/10.1093/hmg/ddr306.

[26] C. Settembre, A. Fraldi, D.L. Medina, A. Ballabio, Signals from the lysosome: a control centre for cellular clearance and energy metabolism, Nat. Rev. Mol. Cell Biol. 14 (2013) 283–296. https://doi.org/10.1038/nrm3565.

[27] R.M. Nisbet, J. Götz, Amyloid-β and Tau in Alzheimer’s Disease: Novel Pathomechanisms and Non-Pharmacological Treatment Strategies, J. Alzheimers Dis. JAD. 64 (2018) S517–S527. https://doi.org/10.3233/JAD-179907.

[28] K.P. Daily, A. Amer, Microglial autophagy-mediated clearance of amyloid-beta plaques is dysfunctional in Alzheimer’s disease mice: Molecular and cell biology/protein clearance/recycling, Alzheimers Dement. 16 (2020). https://doi.org/10.1002/alz.044120.

[29] K. Senkevich, Z. Gan-Or, Autophagy lysosomal pathway dysfunction in Parkinson’s disease; evidence from human genetics, Parkinsonism Relat. Disord. 73 (2020) 60–71. https://doi.org/10.1016/j.parkreldis.2019.11.015.

[30] F. Hopfner, S.H. Mueller, S. Szymczak, O. Junge, L. Tittmann, S. May, K. Lohmann, H. Grallert, W. Lieb, K. Strauch, M. Müller-Nurasyid, K. Berger, B. Schormair, J. Winkelmann, B. Mollenhauer, C. Trenkwalder, W. Maetzler, D. Berg, M. Kasten, C. Klein, G.U. Höglinger, T. Gasser, G. Deuschl, A. Franke, M. Krawczak, A. Dempfle, G. Kuhlenbäumer, Rare Variants in Specific Lysosomal Genes Are Associated With Parkinson’s Disease, Mov. Disord. Off. J. Mov. Disord. Soc. 35 (2020) 1245–1248. https://doi.org/10.1002/mds.28037.

[31] F. Wang, C.-S. Xu, W.-H. Chen, S.-W. Duan, S.-J. Xu, J.-J. Dai, Q.-W. Wang, Identification of Blood-Based Glycolysis Gene Associated with Alzheimer’s Disease by Integrated Bioinformatics Analysis, J. Alzheimers Dis. JAD. 83 (2021) 163–178. https://doi.org/10.3233/JAD-210540.

[32] M. Boldridge, J. Shimabukuro, K. Nakamatsu, C. Won, C. Jansen, H. Turner, L. Wang, Characterization of the C-terminal tail of the Arc protein, PloS One. 15 (2020) e0239870. https://doi.org/10.1371/journal.pone.0239870.

[33] Y.-Z. Chen, M. Matsushita, S. Girirajan, M. Lisowski, E. Sun, Y. Sul, R. Bernier, A. Estes, G. Dawson, N. Minshew, G.D. Shellenberg, E.E. Eichler, M.J. Rieder, D.A. Nickerson, D.W. Tsuang, M.T. Tsuang, E.M. Wijsman, W.H. Raskind, Z. Brkanac, Evidence for involvement of GNB1L in autism, Am. J. Med. Genet. Part B Neuropsychiatr. Genet. Off. Publ. Int. Soc. Psychiatr. Genet. 159B (2012) 61–71. https://doi.org/10.1002/ajmg.b.32002.

[34] N.H. Chapman, R.A. Bernier, S.J. Webb, J. Munson, E.M. Blue, D.-H. Chen, E. Heigham, W.H. Raskind, E.M. Wijsman, Replication of a rare risk haplotype on 1p36.33 for autism spectrum disorder, Hum. Genet. 137 (2018) 807–815. https://doi.org/10.1007/s00439-018-1939-3.

[35] Y.H. Soung, S. Ford, C. Yan, J. Chung, The Role of Arrestin Domain-Containing 3 in Regulating Endocytic Recycling and Extracellular Vesicle Sorting of Integrin β4 in Breast Cancer, Cancers. 10 (2018) E507. https://doi.org/10.3390/cancers10120507.

[36] M.R. Dores, H. Lin, N. J Grimsey, F. Mendez, J. Trejo, The α-arrestin ARRDC3 mediates ALIX ubiquitination and G protein-coupled receptor lysosomal sorting, Mol. Biol. Cell. 26 (2015) 4660–4673. https://doi.org/10.1091/mbc.E15-05-0284.

[37] H. Rosenbrock, M. Desch, O. Kleiner, C. Dorner-Ciossek, B. Schmid, S. Keller, C. Schlecker, V. Moschetti, S. Goetz, K.-H. Liesenfeld, G. Fillon, R. Giovannini, S. Ramael, G. Wunderlich, S. Wind, Evaluation of Pharmacokinetics and Pharmacodynamics of BI 425809, a Novel GlyT1 Inhibitor: Translational Studies, Clin. Transl. Sci. 11 (2018) 616–623. https://doi.org/10.1111/cts.12578.

[38] X. Zhao, L. Huang, Y. Lu, W. Jiang, Y. Song, B. Qiu, D. Tao, Y. Liu, Y. Ma, PIWIL2 interacting with IKK to regulate autophagy and apoptosis in esophageal squamous cell carcinoma, Cell Death Differ. 28 (2021) 1941–1954. https://doi.org/10.1038/s41418-020-00725-4.

[39] S. Zhang, Y. Mao, X. Fan, Inhibition of ghrelin o-acyltransferase attenuated lipotoxicity by inducing autophagy via AMPK-mTOR pathway, Drug Des. Devel. Ther. 12 (2018) 873–885. https://doi.org/10.2147/DDDT.S158985.

[40] I. Tanida, T. Ueno, E. Kominami, LC3 conjugation system in mammalian autophagy, Int. J. Biochem. Cell Biol. 36 (2004) 2503–2518. https://doi.org/10.1016/j.biocel.2004.05.009.

[41] N.S. Beesabathuni, P.S. Shah, Quantitative and temporal measurement of dynamic autophagy rates, Systems Biology, 2021. https://doi.org/10.1101/2021.12.06.471515.

[42] Y. Ogasawara, J. Cheng, T. Tatematsu, M. Uchida, O. Murase, S. Yoshikawa, Y. Ohsaki, T. Fujimoto, Long-term autophagy is sustained by activation of CCTβ3 on lipid droplets, Nat. Commun. 11 (2020) 4480. https://doi.org/10.1038/s41467-020-18153-w.

[43] Y. Chen, M.B. Azad, S.B. Gibson, Superoxide is the major reactive oxygen species regulating autophagy, Cell Death Differ. 16 (2009) 1040–1052. https://doi.org/10.1038/cdd.2009.49.

[44] M. Groh, L.O. Albulescu, A. Cristini, N. Gromak, Senataxin: Genome Guardian at the Interface of Transcription and Neurodegeneration, J. Mol. Biol. 429 (2017) 3181–3195. https://doi.org/10.1016/j.jmb.2016.10.021.

[45] A. Kannan, J. Cuartas, P. Gangwani, D. Branzei, L. Gangwani, Mutation in senataxin alters the mechanism of R-loop resolution in amyotrophic lateral sclerosis 4, Brain J. Neurol. (2022) awab464. https://doi.org/10.1093/brain/awab464.

[46] S.K. Sariki, P.K. Sahu, U. Golla, V. Singh, G.K. Azad, R.S. Tomar, Sen1, the homolog of human Senataxin, is critical for cell survival through regulation of redox homeostasis, mitochondrial function, and the TOR pathway in *Saccharomyces cerevisiae*, FEBS J. 283 (2016) 4056–4083. https://doi.org/10.1111/febs.13917.

[47] C.L. Bennett, A.R. La Spada, SUMOylated Senataxin functions in genome stability, RNA degradation, and stress granule disassembly, and is linked with inherited ataxia and motor neuron disease, Mol. Genet. Genomic Med. 9 (2021). https://doi.org/10.1002/mgg3.1745.

[48] M. Jurga, A.A. Abugable, A.S.H. Goldman, S.F. El-Khamisy, USP11 controls R-loops by regulating senataxin proteostasis, Nat. Commun. 12 (2021) 5156. https://doi.org/10.1038/s41467-021-25459-w.

[49] P. Richard, S. Feng, Y.-L. Tsai, W. Li, P. Rinchetti, U. Muhith, J. Irizarry-Cole, K. Stolz, L.A. Sanz, S. Hartono, M. Hoque, S. Tadesse, H. Seitz, F. Lotti, M. Hirano, F. Chédin, B. Tian, J.L. Manley, SETX (senataxin), the helicase mutated in AOA2 and ALS4, functions in autophagy regulation, Autophagy. 17 (2021) 1889–1906. https://doi.org/10.1080/15548627.2020.1796292.

[50] C.L. Bennett, B.L. Sopher, A.R. La Spada, Tight expression regulation of senataxin, linked to motor neuron disease and ataxia, is required to avert cell-cycle block and nucleolus disassembly, Heliyon. 6 (2020) e04165. https://doi.org/10.1016/j.heliyon.2020.e04165.

[51] N. Zielke, K.J. Kim, V. Tran, S.T. Shibutani, M.-J. Bravo, S. Nagarajan, M. van Straaten, B. Woods, G. von Dassow, C. Rottig, C.F. Lehner, S.S. Grewal, R.J. Duronio, B.A. Edgar, Control of Drosophila endocycles by E2F and CRL4(CDT2), Nature. 480 (2011) 123–127. https://doi.org/10.1038/nature10579.

[52] M. Xiong, W. Hu, Y. Tan, H. Yu, Q. Zhang, C. Zhao, Y. Yi, Y. Wang, Y. Wu, M. Wu, Transcription Factor E2F1 Knockout Promotes Mice White Adipose Tissue Browning Through Autophagy Inhibition, Front. Physiol. 12 (2021) 748040. https://doi.org/10.3389/fphys.2021.748040.

[53] W. Kim, Y.-G. Jang, J. Yang, J. Chung, Spatial Activation of TORC1 Is Regulated by Hedgehog and E2F1 Signaling in the Drosophila Eye, Dev. Cell. 42 (2017) 363-375.e4. https://doi.org/10.1016/j.devcel.2017.07.020.

[54] S. Polager, M. Ofir, D. Ginsberg, E2F1 regulates autophagy and the transcription of autophagy genes, Oncogene. 27 (2008) 4860–4864. https://doi.org/10.1038/onc.2008.117.

[55] K. Motonaga, M. Itoh, A. Hirayama, S. Hirano, L.E. Becker, Y. Goto, S. Takashima, Up-regulation of E2F-1 in Down’s syndrome brain exhibiting neuropathological features of Alzheimer-type dementia, Brain Res. 905 (2001) 250–253. https://doi.org/10.1016/S0006-8993(01)02535-5.

[56] S. Gehrke, Y. Imai, N. Sokol, B. Lu, Pathogenic LRRK2 negatively regulates microRNA-mediated translational repression, Nature. 466 (2010) 637–641. https://doi.org/10.1038/nature09191.

[57] G.U. Hoglinger, J.J. Breunig, C. Depboylu, C. Rouaux, P.P. Michel, D. Alvarez-Fischer, A.-L. Boutillier, J. DeGregori, W.H. Oertel, P. Rakic, E.C. Hirsch, S. Hunot, The pRb/E2F cell-cycle pathway mediates cell death in Parkinson’s disease, Proc. Natl. Acad. Sci. 104 (2007) 3585–3590. https://doi.org/10.1073/pnas.0611671104.

[58] D.S. Castillo, A. Campalans, L.M. Belluscio, A.L. Carcagno, J.P. Radicella, E.T. Cánepa, N. Pregi, E2F1 and E2F2 induction in response to DNA damage preserves genomic stability in neuronal cells, Cell Cycle. 14 (2015) 1300–1314. https://doi.org/10.4161/15384101.2014.985031.

[59] L.K. Hein, P.M. Apaja, K. Hattersley, R.H. Grose, J. Xie, C.G. Proud, T.J. Sargeant, A novel fluorescent probe reveals starvation controls the commitment of amyloid precursor protein to the lysosome, Biochim. Biophys. Acta BBA - Mol. Cell Res. 1864 (2017) 1554–1565. https://doi.org/10.1016/j.bbamcr.2017.06.011.

[60] M. Martin, Cutadapt removes adapter sequences from high-throughput sequencing reads, EMBnet.Journal. 17 (2011) 10. https://doi.org/10.14806/ej.17.1.200.

[61] K.L. Howe, B. Contreras-Moreira, N. De Silva, G. Maslen, W. Akanni, J. Allen, J. Alvarez-Jarreta, M. Barba, D.M. Bolser, L. Cambell, M. Carbajo, M. Chakiachvili, M. Christensen, C. Cummins, A. Cuzick, P. Davis, S. Fexova, A. Gall, N. George, L. Gil, P. Gupta, K.E. Hammond-Kosack, E. Haskell, S.E. Hunt, P. Jaiswal, S.H. Janacek, P.J. Kersey, N. Langridge, U. Maheswari, T. Maurel, M.D. McDowall, B. Moore, M. Muffato, G. Naamati, S. Naithani, A. Olson, I. Papatheodorou, M. Patricio, M. Paulini, H. Pedro, E. Perry, J. Preece, M. Rosello, M. Russell, V. Sitnik, D.M. Staines, J. Stein, M.K. Tello-Ruiz, S.J. Trevanion, M. Urban, S. Wei, D. Ware, G. Williams, A.D. Yates, P. Flicek, Ensembl Genomes 2020—enabling non-vertebrate genomic research, Nucleic Acids Res. 48 (2020) D689–D695. https://doi.org/10.1093/nar/gkz890.

[62] A. Dobin, C.A. Davis, F. Schlesinger, J. Drenkow, C. Zaleski, S. Jha, P. Batut, M. Chaisson, T.R. Gingeras, STAR: ultrafast universal RNA-seq aligner, Bioinforma. Oxf. Engl. 29 (2013) 15–21. https://doi.org/10.1093/bioinformatics/bts635.

[63] Y. Liao, G.K. Smyth, W. Shi, featureCounts: an efficient general purpose program for assigning sequence reads to genomic features, Bioinforma. Oxf. Engl. 30 (2014) 923–930. https://doi.org/10.1093/bioinformatics/btt656.

[64] C.M. Ward, T.-H. To, S.M. Pederson, ngsReports: a Bioconductor package for managing FastQC reports and other NGS related log files, Bioinforma. Oxf. Engl. 36 (2020) 2587–2588. https://doi.org/10.1093/bioinformatics/btz937.

[65] K.D. Hansen, R.A. Irizarry, Z. Wu, Removing technical variability in RNA-seq data using conditional quantile normalization, Biostatistics. 13 (2012) 204–216. https://doi.org/10.1093/biostatistics/kxr054.

[66] S. Lund, Detecting Differential Expression in RNA-Sequence Data Using Quasi-Likelihood with Shrunken Dispersion Estimates, (2012). https://tsapps.nist.gov/publication/get_pdf.cfm?pub_id=911007.

[67] M.D. Robinson, D.J. McCarthy, G.K. Smyth, edgeR: a Bioconductor package for differential expression analysis of digital gene expression data, Bioinforma. Oxf. Engl. 26 (2010) 139–140. https://doi.org/10.1093/bioinformatics/btp616.

[68] D.J. McCarthy, G.K. Smyth, Testing significance relative to a fold-change threshold is a TREAT, Bioinformatics. 25 (2009) 765–771. https://doi.org/10.1093/bioinformatics/btp053.

[69] A.L. Tarca, S. Draghici, P. Khatri, S.S. Hassan, P. Mittal, J. Kim, C.J. Kim, J.P. Kusanovic, R. Romero, A novel signaling pathway impact analysis, Bioinformatics. 25 (2009) 75–82. https://doi.org/10.1093/bioinformatics/btn577.

[70] M. Kanehisa, S. Goto, KEGG: kyoto encyclopedia of genes and genomes, Nucleic Acids Res. 28 (2000) 27–30. https://doi.org/10.1093/nar/28.1.27.

[71] G. Sales, E. Calura, C. Romualdi, metaGraphite-a new layer of pathway annotation to get metabolite networks, Bioinforma. Oxf. Engl. 35 (2019) 1258–1260. https://doi.org/10.1093/bioinformatics/bty719.

[72] I. Yevshin, R. Sharipov, S. Kolmykov, Y. Kondrakhin, F. Kolpakov, GTRD: a database on gene transcription regulation-2019 update, Nucleic Acids Res. 47 (2019) D100–D105. https://doi.org/10.1093/nar/gky1128.

[73] A. Liberzon, A. Subramanian, R. Pinchback, H. Thorvaldsdóttir, P. Tamayo, J.P. Mesirov, Molecular signatures database (MSigDB) 3.0, Bioinforma. Oxf. Engl. 27 (2011) 1739–1740. https://doi.org/10.1093/bioinformatics/btr260.

