## Supplementary material for "Transcriptional targets of senataxin and E2 promoter binding factors are associated with neuro-degenerative pathways during increased autophagic flux": manuscript_v8.docx

### Competing Interests

The authors declare no competing interests.

### Figure 1

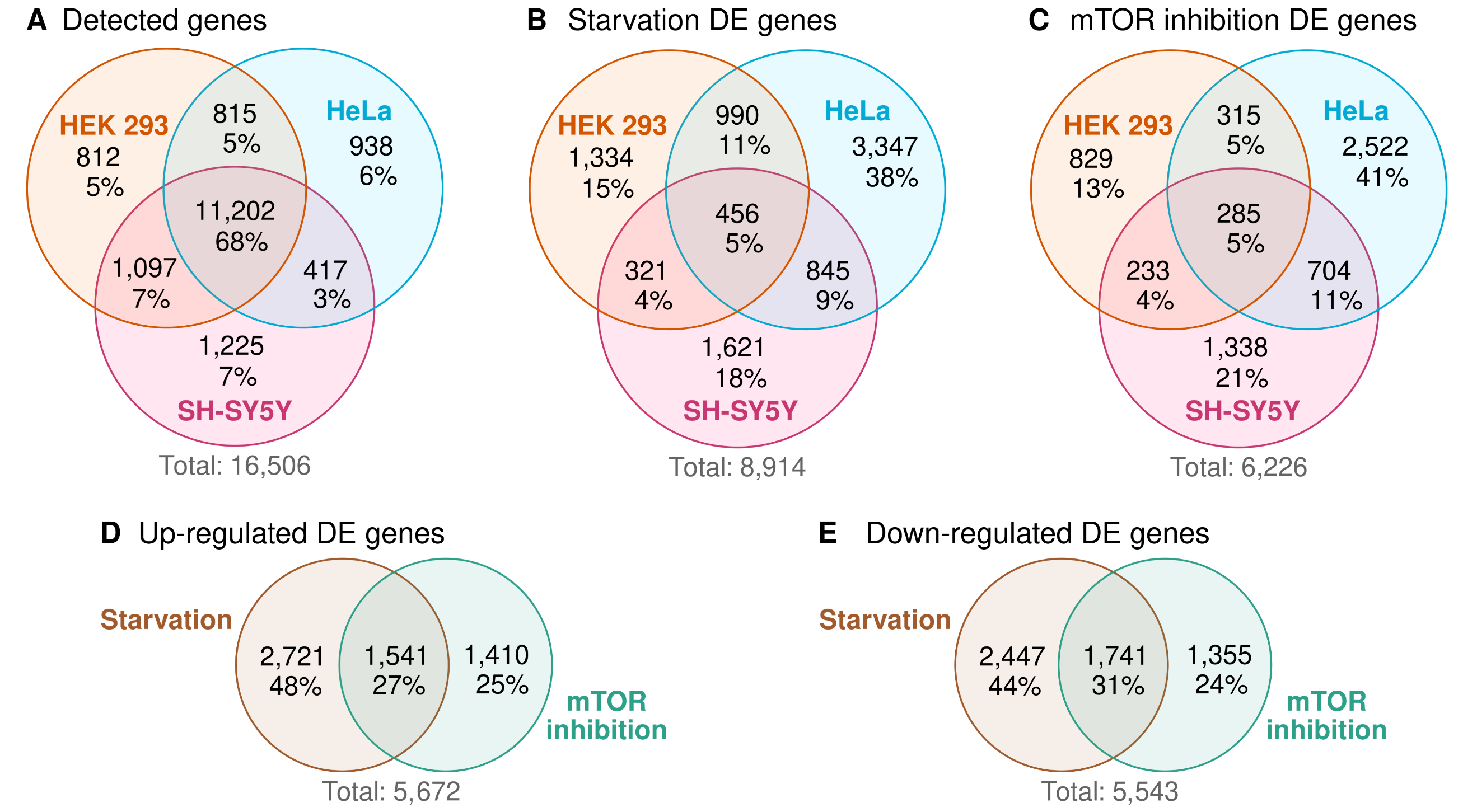

Overview of differentially expressed (DE) genes. **A**) Genes were considered detectable if there were >1.5 counts per million in >3 samples out of all samples from the same cell line. **B**) Genes that were DE between starved and control samples in at least one cell line. **C**) Genes that were DE between mTOR inhibited and control samples in at least one cell line. **D**) We collected DE genes associated with starvation or mTOR inhibition that were up-regulated in at least one cell line (inconsistent DE genes that were significantly up-regulated in one cell line but significantly down-regulated in another were excluded). **E**) Down-regulated DE genes associated with starvation or mTOR inhibition.

### Figure 2

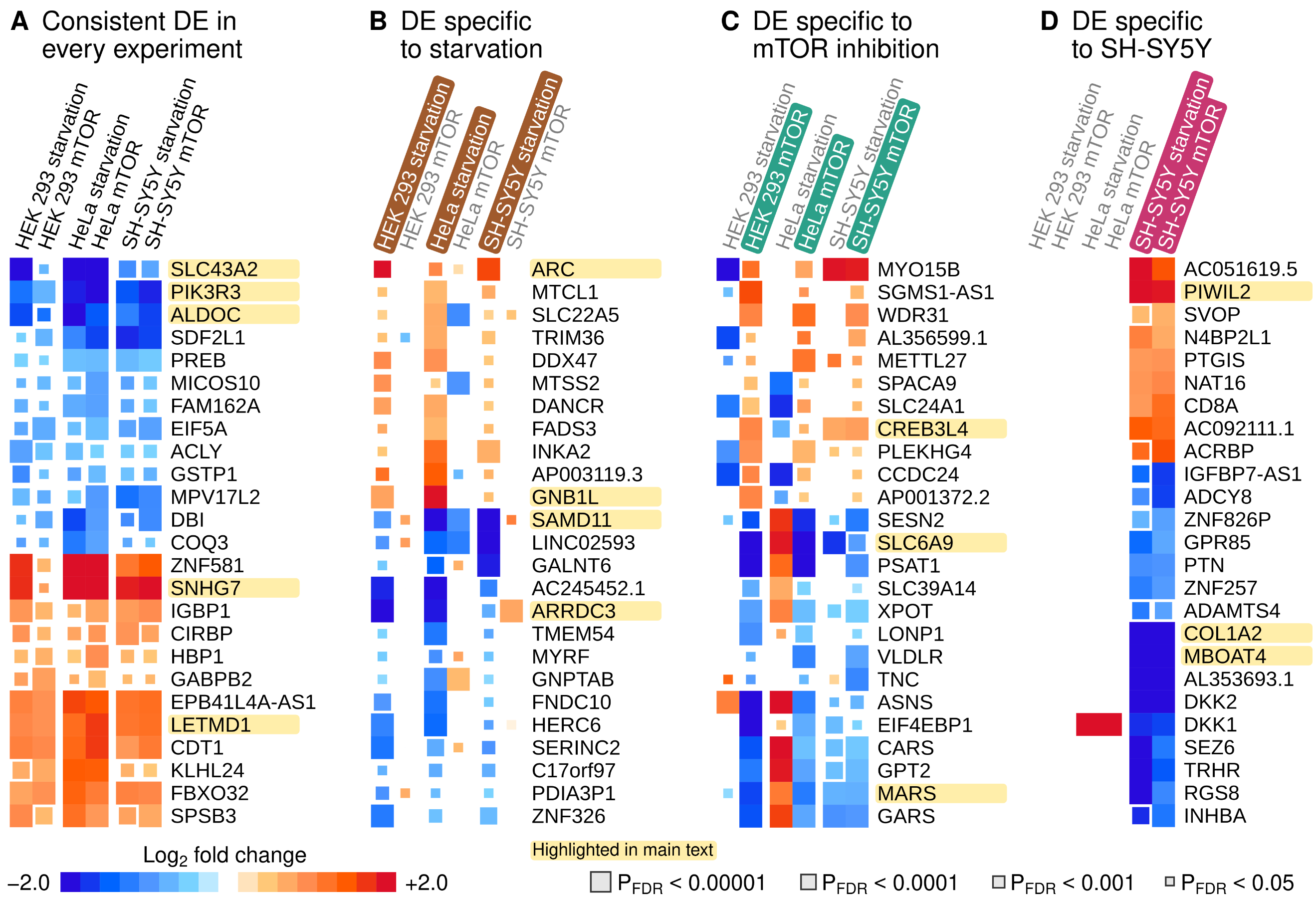

Top 25 differentially expressed (DE) genes based on the maximum FDR rule. Genes mentioned in the main text are highlighted for easier visual localization. **A**) Genes were sorted according to the maximum FDR-adjusted P-value across six experiments. Discordant genes that were significantly (P_FDR_ < 5%) up-regulated in one and down-regulated in another experiment were excluded. **B**) Genes were sorted according to the maximum P_FDR_ across all starvation experiments. We also required that all starvation responses were directionally concordant and that the mean log_2_ fold change across mTOR experiments was in the opposite direction. **C**) Genes were sorted according to the maximum P_FDR_ across all mTOR inhibition experiments. We required that all mTOR inhibition responses were directionally concordant and that the mean log_2_ fold change across starvation experiments was in the opposite direction. **D**) Genes were sorted according to the maximum P_FDR_ across responses in the SH-SY5Y cells. Missing signals were set to zero log_2_ fold change in other cell lines. We also required that the mean log_2_ fold changes in other cells were in the opposite direction to SH-SY5Y responses.

### Figure 3

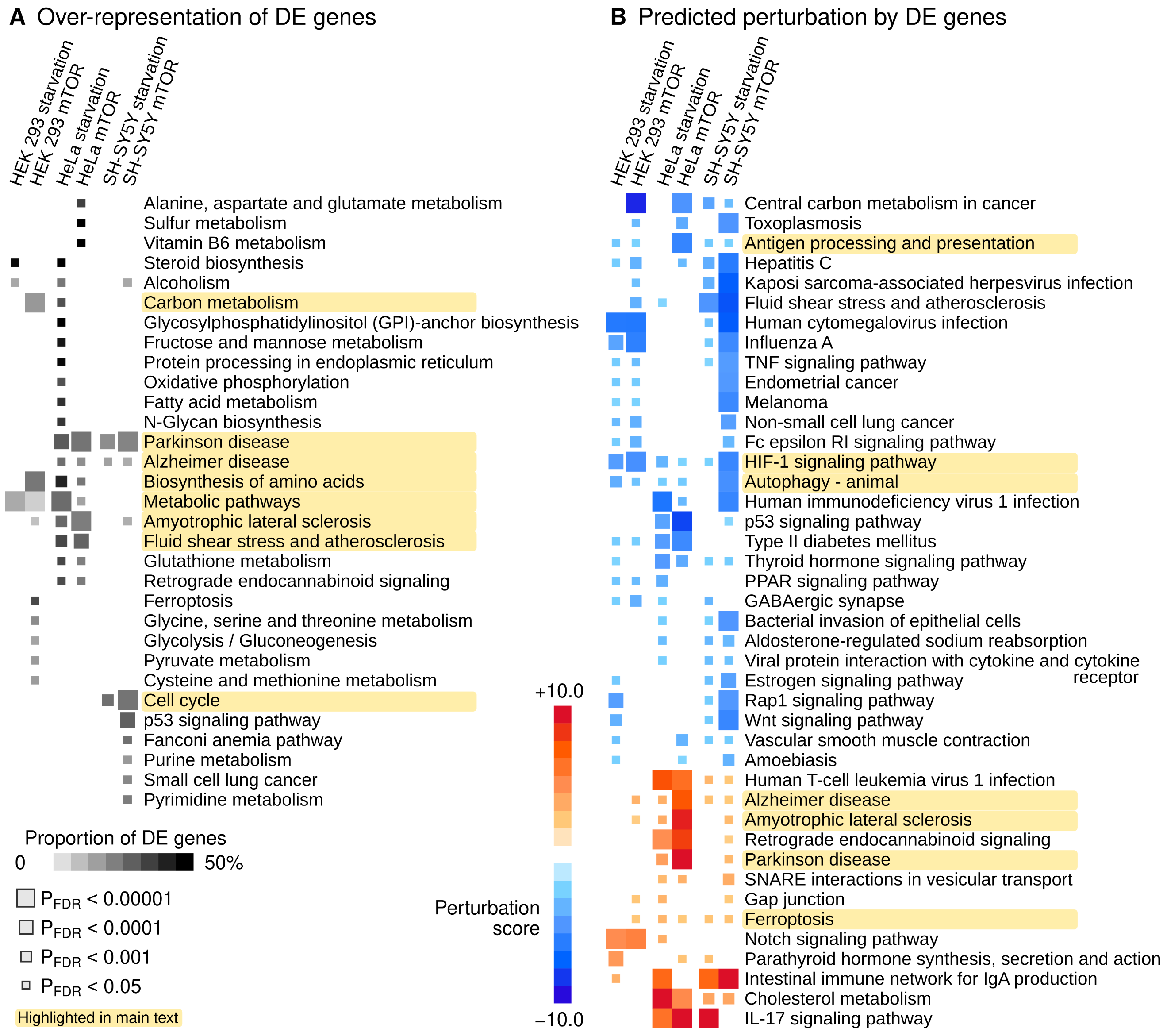

Enrichment of differentially expressed genes in the Kyoto Encyclopedia of Genes and Genomes pathway repository. **A**) Over-representation analysis of DE genes. Pathways that produced a significant signal (P_FDR_ < 0.05) in at least one experiment are shown. **B**) Normalized perturbation scores from Signaling Pathway Impact Analysis. A negative (positive) score implies that the aggregate impact of DE genes is likely to decrease (increase) the activity of a pathway. Pathways that were directionally concordant (all significant signals in the same direction) and that produced at least three significant signals (P_FDR_ < 0.05) are included.

### Figure 4

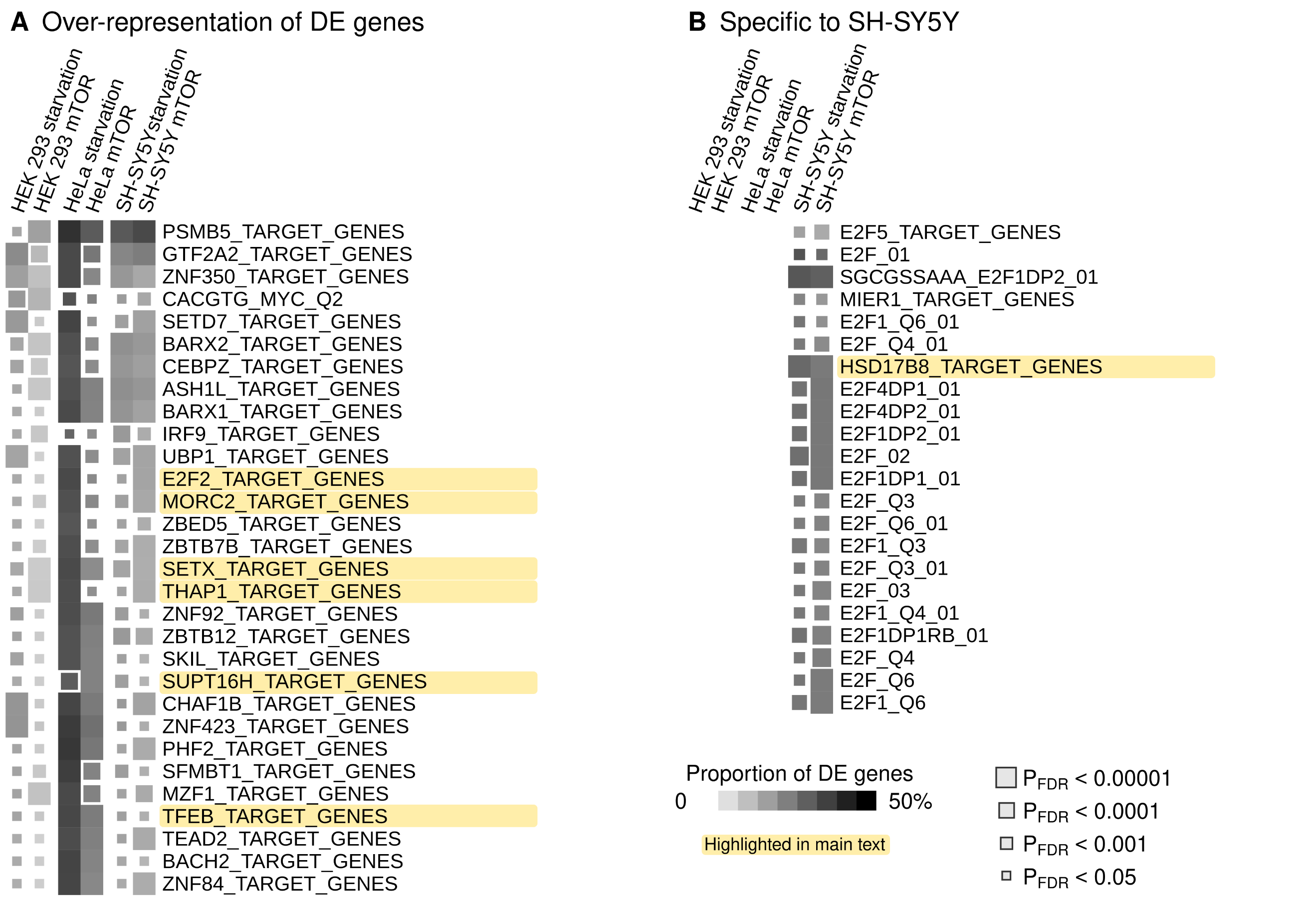

Enrichment of differentially expressed genes in transcription factor target (TFT) sets. **A**) Over-representation analysis of DE genes. Pathways that produced a significant signal (P_FDR_ < 0.05) in at least one experiment are shown. **B**) DE enrichment within TFT sets in SH-SY5Y cells but not in other cells.

### Figure 5

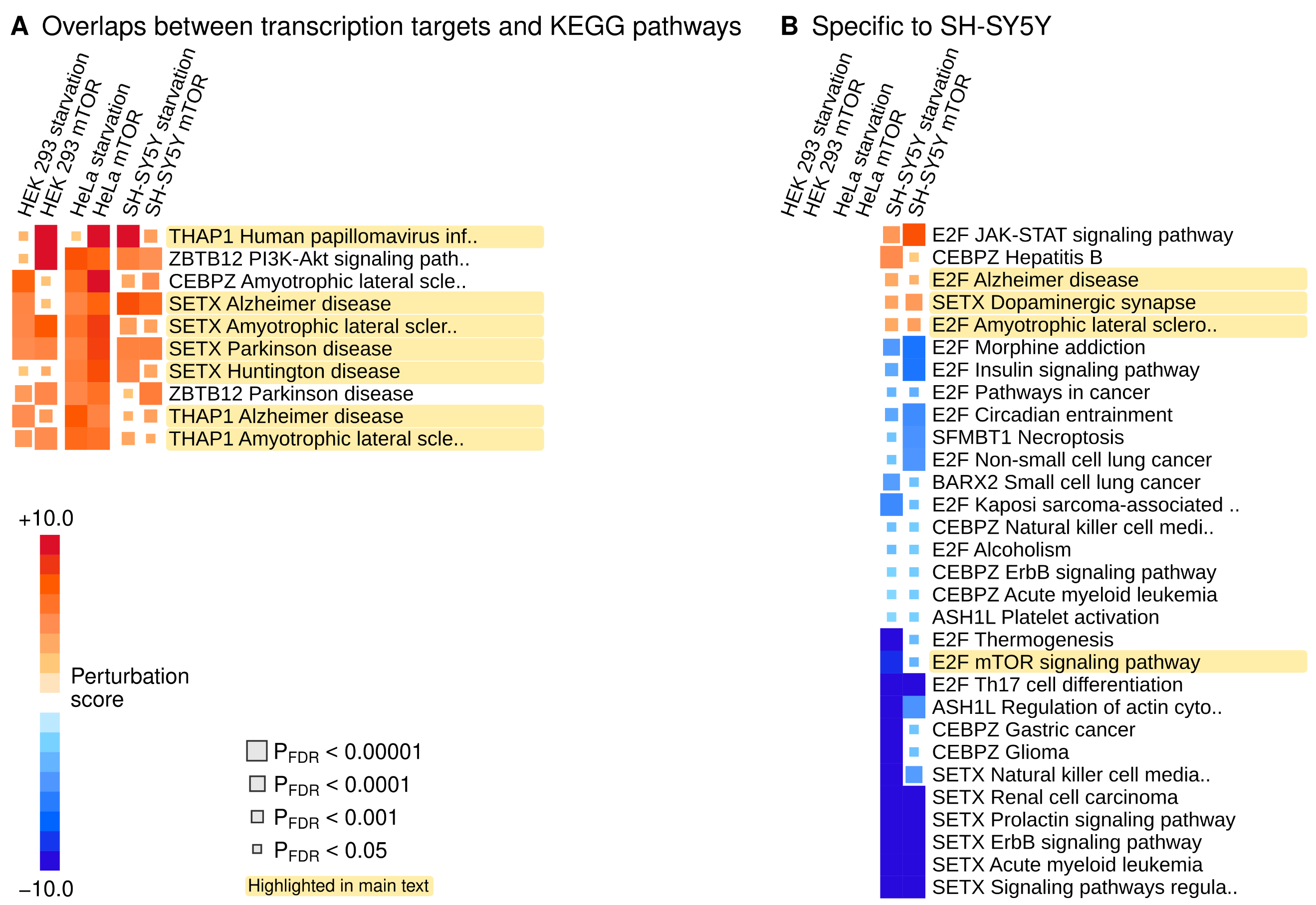

Combined perturbation analysis of canonical pathways and TFT sets. First, we identified DE genes that were shared between a KEGG pathway and TFT sets. Then, we used Signaling Pathway Impact analyses to test if the shared genes would impact the activity of the KEGG pathway. Therefore, the perturbation scores are predictions on the potential regulatory effects differentially expressed transcription factor target genes will have on canonical pathways. **A**) TFT-pathway pairs that showed directionally consistent and significant (P_FDR_ < 0.05) perturbation scores across every experiment. **B**) TFT-pathway pairs that showed directionally consistent and significant (P_FDR_ < 0.05) perturbation scores in the two experiments on SH-SY5Y cells but no significant signals in other cells.
