## Supplementary material for "Transcriptional targets of senataxin and E2 promoter binding factors are associated with neuro-degenerative pathways during increased autophagic flux": supplement_material_v8.docx

### Supplementary Figure S1

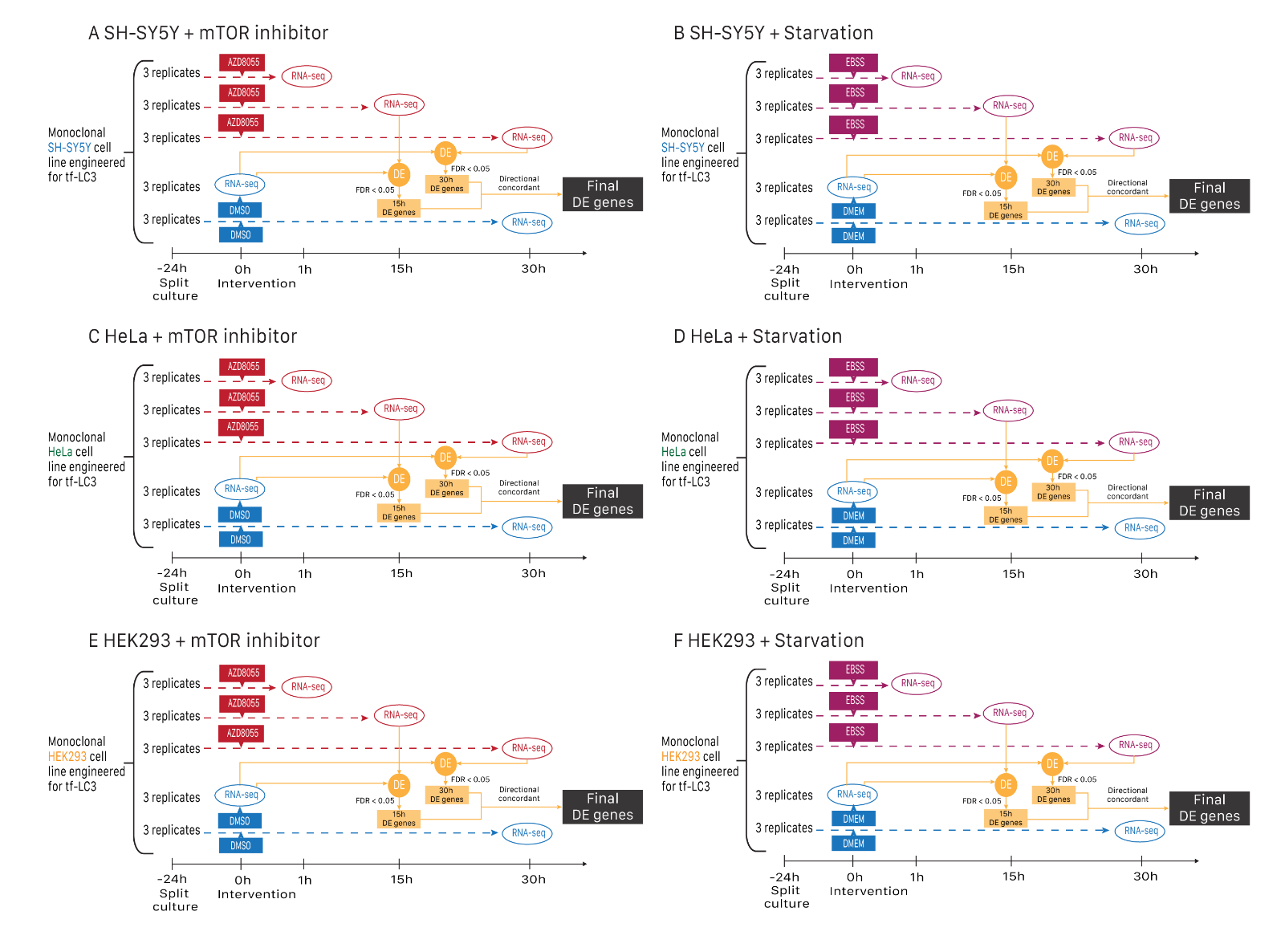

Schematic illustration of how differentially expressed (DE) genes were determined for each combination of cell line and treatment. Inhibition of the mTOR complexes was achieved by the chemical combound AZD8055. Starvation was induced by applying Earle's Balanced Salt Solution (EBSS) as culture media.

### Supplementary Figure S2

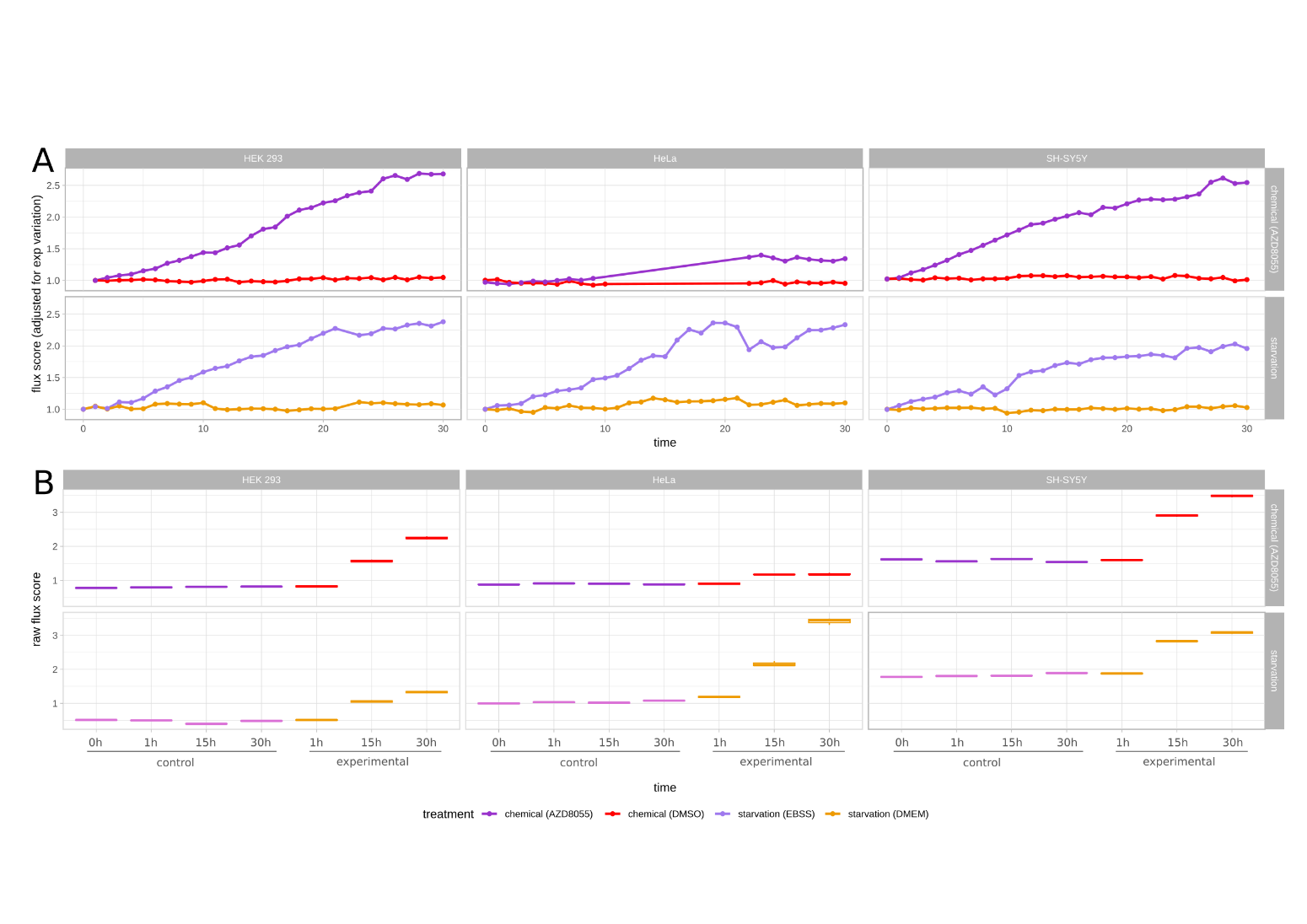
Assessment of autophagic flux using the tf-LC3 assay. The flux score was defined as the ratio between the red and green fluorophores (see Methods). **A**) Monoclonal cultures of three cell lines that were subjected to starvation (EBSS) and chemical mTOR inhibition (AZD8055). Samples were obtained every hour to determine the time curves of autophagy responses. **B**) Measurements of autophagic flux from the samples that were sent to RNA-sequencing. Interquartile box plots of three replicates are shown, although variation was so low that the boxes got flattened for most time points.

### Supplementary Figure S3

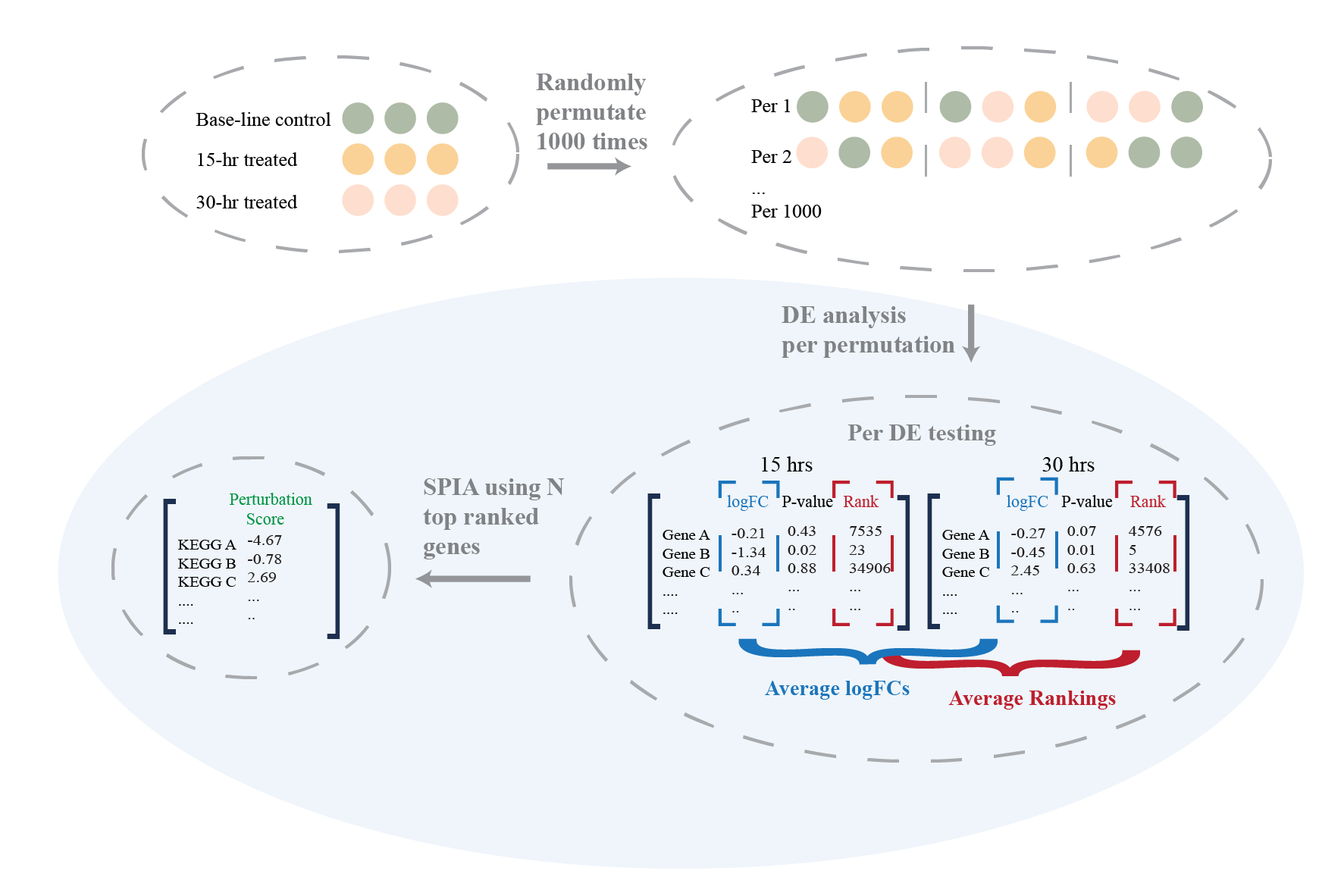

In Signaling Pathway Impact Analysis (SPIA), the probability of obtaining the observed perturbation score (PS) is calculated through a bootstrap approach, where random genes in the same numbers as DE genes inputted would be assigned to random locations of pathways and receive logFCs randomly sampled from the range of logFC as observed within DE genes. The underlying assumption of this approach is that genes are statistically independent, while the primary purpose of gene-sets was to group co-regulated genes, thus challenging the assumption. Moreover, assigning genes randomly breaks the gene-gene correlations, which could result in over-estimated statistical significance [Dørum, G., et al., Rotation testing in gene set enrichment analysis for small direct comparison experiments. Stat Appl Genet Mol Biol, 2009. 8: p. Article34. Efron, B. and R. Tibshirani, On testing the significance of sets of genes. The annals of applied statistics, 2007. 1(1): p. 107-129.]. Therefore, we developed an alternative significance testing method that permutes samples instead to simulate null distributions of KEGG pathways PSs.

To simulate null distributions of PSs, we firstly generated permuted logFCs by shuffling the 9 samples within each condition (3 base-line control, 3 15 h-treated and 3 30 h treated) and assigning the first 3 samples of each permutation to be base-line controls, middle 3 to be 15 h-treated and last 3 to be 30-hr treated samples. 1000 permutations were randomly sampled from the total 362,880 possible permutations and differential expression analyses were performed 1000 times following methods described in the ‘Differential Expression Analysis” section. To reduce computation time, dispersion estimation required in the edgeR workflow was only performed once, assuming that all samples were derived from the same group.

For each round of DE testing, genes were ranked by p-values at both time points, and the final rankings were derived through ranking the mean of two individual rankings. The same number of top-ranked genes as the number of DEGs defined under each condition were kept and used as the permuted DEGs. Again, average logFCs between 15 h and 30 h were taken for each DE testing to derive a single estimate of FCs, thus giving rise to 1000 sets of permuted DEGs with permuted logFCs for each condition.

SPIA’s net perturbation accumulation algorithm was then applied 1000 times utilising the permutated DEGs and their logFCs to derive the null distribution of perturbation scores for each KEGG pathway, from which the mean standard deviation (MAD) of each pathway was calculated. As the default of mad function in the *stats* package, the actual value outputted was constant × MAD, where the constant was defaulted to be 1.4826 to approximate standard deviation. Statistical significance of observed PS was then defined by firstly calculating the corresponding robust z-scores:$\frac{(tA-median)}{scaled MAD}$. The robust z-scores were then converted to 2-sided p-values and multiple testing consideration was accounted for through the Benjamini-Hochberg method. KEGG pathways with perturbation FDR smaller than 0.05 were deemed to be significantly perturbed.
